# Kcs1-dependent inositol pyrophosphates promote DNA double-strand break repair via Pho80-Pho85 in budding yeast

**DOI:** 10.64898/2026.10.07.751851

**Authors:** Giovanna Marques Panessa, Paulo Pereira Sforsin, Mateusz Maciej Wagner, Giuliana Monteiro, Marcus Bustamante Smolka, José Renato Rosa Cussiol

## Abstract

The cleavage of membrane-bound phosphoinositides produces inositol polyphosphates (IPs) and pyrophosphates (PP-IPs), small water-soluble signaling molecules that are highly conserved in eukaryotes and play a critical role in cell metabolism. Loss of the inositol pyrophosphate kinase (PP-IPk) Kcs1 and its mammalian ortholog IP6K1 was previously shown to cause hypersensitivity to genotoxic stress and reduced homologous recombination (HR) repair. To identify the mechanism linking PP-IPs to DNA double-strand break (DSB) repair, we combined quantitative proteomics with biochemical, genetic, and cell-cycle analyses in budding yeast. We found that loss of Kcs1, but not the PP-IPk Vip1, caused increased sensitivity to the radiomimetic drug zeocin. *kcs1Δ* cells displayed persistent Rad53 signaling and elevated γ-H2A levels, remaining arrested in G1/S during treatment and accumulating in G2/M for hours after damage removal. Unexpectedly, Kcs1-dependent PP-IPs promoted DSB repair independently of the core HR machinery and of cell-cycle stage. Quantitative proteomics revealed increased levels of PHO pathway proteins in zeocin-treated *kcs1Δ* cells, and deletion of the CDK inhibitor *PHO81* substantially rescued zeocin hypersensitivity, indicating that constitutive inhibition of the Pho80-Pho85 cyclin-dependent kinase underlies the DNA repair defect caused by Kcs1 loss. This function of Pho80-Pho85 was mechanistically distinct from its canonical role in phosphate homeostasis, and partially independent of its connections to Ku80 regulation. Together, our findings identify Kcs1-dependent PP-IP signaling as a new upstream input controlling Pho80-Pho85 activity during the DNA damage response, establishing an HR-independent axis for genome maintenance and linking PP-IP metabolism to the DNA damage response in eukaryotic cells.

## Introduction

In eukaryotes, the inositol polyphosphate kinases (IPKs) are responsible for catalyzing the transfer of a phosphate group from ATP to specific hydroxyl groups on the inositol ring leading to the formation of inositol polyphosphates (IPs). These molecules are precursors for the formation of inositol pyrophosphates (PP-IPs) which carry a diphosphate group at either the 1- or 5-position of the inositol ring (1). PP-IP synthesis is dependent on the activity of the PP-IP kinases Kcs1 (ortholog of IP6K1-3 in mammals) and Vip1 (ortholog of PPIP5K1 and PPIP5K2 in mammals) while inositol pyrophosphatases such as Ddp1 and Siw14 can rapidly remove a pyrophosphate from inositol (2–5) (Fig. 1).

**Figure 1.**
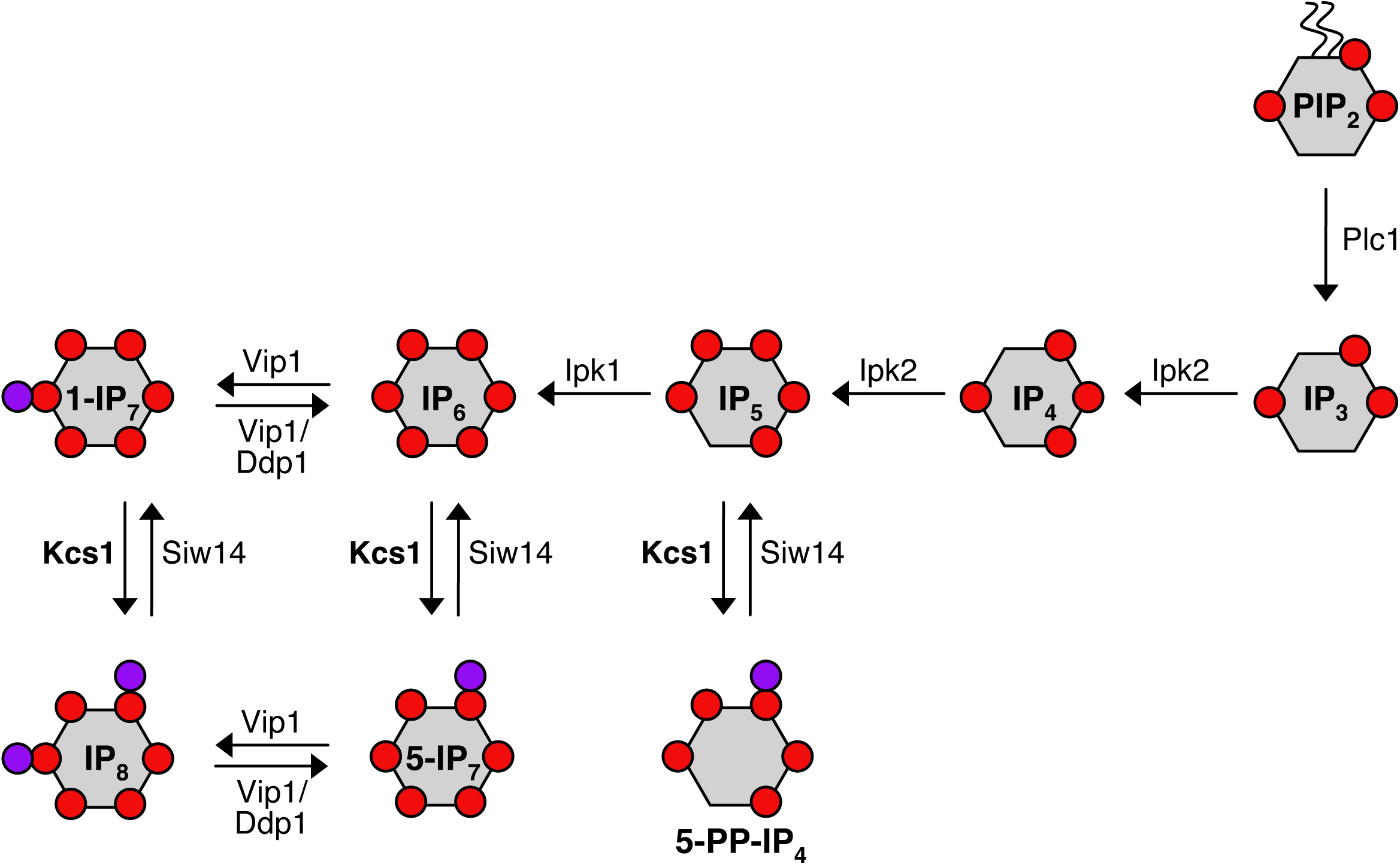
The inositol polyphosphate biosynthesis pathway in budding yeast. Phospholipase C (Plc1) hydrolyzes phosphatidylinositol 4,5-bisphosphate (PIP2) to generate the signaling molecules inositol 1,4,5-trisphosphate (IP3) and 1,2-diacylglycerol (DAG). IP3 is sequentially phosphorylated to IP4 and IP5 by Ipk2, followed by phosphorylation of IP5 to IP6 by Ipk1. Kcs1 generates 5-PP-IP₄ and 5-PP-IP₅ (5-IP₇), whereas Vip1 generates 1-PP-IP₅ (1-IP₇). Further phosphorylation of IP₇ species generates 1,5-IP₈. Siw14, Ddp1, and Vip1 contribute to PP-IP turnover through their inositol pyrophosphatase activities. DAG, 1,2-diacylglycerol, is omitted for clarity.

Because of their small size, water solubility, high-energy pyrophosphate bonds, and rapid turnover, PP-IPs function as powerful signaling molecules in eukaryotes that modulate multiple cellular and metabolic pathways (6–11). PP-IPs exert their biological functions through the regulation of protein function, either by transient interactions with protein domains or through non-enzymatic pyrophosphorylation (12–16). Studies in yeast and mammalian cell lines have established several functions for PP-IPs, such as the regulation of gene expression (17), phosphate metabolism (3, 18, 19), telomere homeostasis (20–22), endocytic trafficking (23), environmental stress response (24–26), DNA replication and repair (27–30), among others. In pathogenic fungi, PP-IPs also contribute to virulence by regulating phosphate metabolism in *Cryptococcus neoformans* (31, 32).

One of the best-characterized effector mechanisms of PP-IP signaling in eukaryotes is the control of phosphate homeostasis through binding to SPX domains which are present in most components of the phosphate-signaling machinery from yeast to plants and humans (33). In budding yeast, this mechanism converges on the cyclin-dependent kinase (CDK) complex Pho80-Pho85. Under phosphate-replete conditions, Pho80-Pho85 phosphorylates the transcription factor Pho4, promoting its nuclear export and keeping the phosphate-responsive PHO regulon repressed (34, 35). Kinase activity is set by Pho81, a CDK inhibitor (CKI) that remains constitutively associated with Pho80-Pho85 and that carries an N-terminal SPX domain (36). Current evidence indicates that 1,5-IP8, generated by the sequential action of Kcs1 and Vip1 (Fig. 1), accumulates when phosphate is abundant and, by binding the SPX domain of Pho81, keeps the inhibitor silent and the kinase active. Upon phosphate starvation, 1,5-IP8 declines, Pho81 inhibits Pho80-Pho85, Pho4 accumulates in the nucleus and the PHO regulon is induced (37). Because *PHO81* is itself a Pho4 target gene (38), inhibition of the kinase also raises Pho81 levels, generating a feed-forward loop that reinforces the inhibited state. Cells lacking Kcs1, which cannot produce 5-IP7 and therefore fail to accumulate 1,5-IP8, phenocopy phosphate starvation: Pho80-Pho85 is constitutively inhibited by Pho81 even in phosphate-replete medium (37, 39). Importantly, Pho85 is a pleiotropic CDK that associates with ten different cyclins and phosphorylates substrates well beyond the PHO regulon (40), so a constitutive block of Pho80-Pho85 activity is expected to have consequences that are not restricted to phosphate metabolism.

The emerging role of PP-IPs in cancer further underscores their importance in cellular homeostasis. IP6K1 and IP6K2 have been implicated in cancer cell invasion and proliferation, while PPIP5K1 and PPIP5K2 have been shown to promote tumor growth through modulation of p53- and Akt/mTOR-dependent pathways (as reviewed by (41)). However, these effects appear context-dependent, as deletion of *IP6K1* or *IP6K2* can either suppress or enhance tumor growth depending on the experimental setting. This observation may reflect, at least in part, differences in how PP-IP metabolism intersects with genotoxic stress responses and DNA repair capacity across cellular contexts. Consistent with this idea, a growing body of mechanistic evidence directly links IPs and PP-IPs to DNA repair. Inositol hexakisphosphate (IP6) binds to the mammalian KU70/KU80 heterodimer and stimulates non-homologous end joining (NHEJ) repair (28, 42, 43). In addition, IPMK (the mammalian ortholog of yeast Ipk2) has been proposed to promote the recruitment of ATR to sites of DNA damage through the synthesis of a nuclear pool of phosphoinositides (44). In mammalian cells, perturbation of PP-IP metabolism affects the cellular response to DNA damage. IP6K1-derived 5-PP-IP5, for instance, is required for efficient recovery from DNA damage, as IP6K1-deficient mouse embryonic fibroblasts (MEFs) display impaired completion of HR repair that can be rescued by catalytically active, but not kinase-inactive, IP6K1 (29). Moreover, PPIP5K2 was shown to promote HR by translocating to the nucleus and recruiting RPA70 to sites of DNA damage, suggesting that PP-IP metabolic enzymes can influence DNA repair through both their enzymatic products and protein-mediated mechanisms (45). In budding yeast, deletion of Kcs1 was shown to restore normal levels of HR in a hyper-recombination strain, suggesting that Kcs1-dependent PP-IPs modulate HR through an unknown mechanism (30). Subsequent studies demonstrated that Kcs1 is required for full resistance to the radiomimetic agent phleomycin (24). We have previously demonstrated that overexpression of Kcs1 leads to hypersensitivity to methyl methanesulfonate (MMS), indicating that precise modulation of intracellular Kcs1-dependent PP-IP levels is critical for an appropriate genotoxic stress response (46). These findings support the notion that inositol phosphate metabolism, and PP-IPs in particular, are important for the DNA damage response (DDR) in eukaryotic cells. However, the molecular mechanisms by which PP-IPs regulate the DDR remain poorly understood.

In this study, we investigated the role of inositol polyphosphate metabolism in the DDR in budding yeast. We show that Kcs1-dependent PP-IPs promote efficient repair of DSBs, and that loss of Kcs1 results in persistent DNA damage signaling and impaired recovery following genotoxic stress induced by the radiomimetic drug zeocin. Moreover, DSB induction by the expression of an HO endonuclease leads to increased γ-H2A levels and loss of viability in *kcs1Δ* cells. Through proteomic, genetic, cell-cycle, and biochemical analyses, we demonstrate that the role of Kcs1 in DSB repair is independent of the core HR machinery and identify the Pho81-Pho80-Pho85 module as a key downstream effector of Kcs1-dependent PP-IP signaling, with loss of Kcs1 causing constitutive, Pho81-mediated inhibition of the kinase. Importantly, this function of Pho80-Pho85 in DSB repair is separable from its canonical role in phosphate homeostasis, as neither the accompanying hyperactivation of the PHO regulon nor the depletion of inorganic polyphosphate accounts for the repair defect. Our findings further reveal an unexpected contribution of Ku80 to the DNA repair defects associated with loss of Kcs1, suggesting a functional link between PP-IP signaling and Ku80-dependent processes during the response to DSBs. Our results uncover a previously unrecognized role for Kcs1-dependent PP-IPs in DSB repair and provide new insights into how inositol phosphate signaling interfaces with genome maintenance.

## Results

### Kcs1-dependent PP-IPs are important for genotoxic stress response

To determine the role of inositol polyphosphate metabolism in the DNA damage response (DDR), we examined the sensitivity of strains lacking the IP kinases Ipk1 (*ipk1Δ*) and Ipk2 (*ipk2Δ*) or the PP-IP kinases Vip1 (*vip1Δ*) and Kcs1 (*kcs1Δ*) to genotoxic stress. Deletion of *IPK2* and *KCS1*, but not *IPK1* or *VIP1*, resulted in pronounced sensitivity to the radiomimetic drug zeocin (Fig. 2*A* and *B*). *kcs1Δ* cells also displayed mild sensitivity to hydroxyurea (HU), but not to MMS or H2O2 (Fig. S1). Genetic interaction analyses showed that the *kcs1Δvip1Δ* double mutant was slightly more zeocin-sensitive than *kcs1Δ* alone (Fig. 2*C*), indicating at most a minor contribution of Vip1-dependent species. However, the resistance of *vip1Δ* single mutants to zeocin argues that Vip1 is not the PP-IP kinase required for zeocin resistance. On the other hand, deletion of *IPK2* partially restored zeocin resistance to *kcs1Δ* cells, and conversely deletion of *KCS1* substantially improved the zeocin survival of *ipk2Δ* cells despite comparable growth of the two strains on drug-free medium (Fig. 2*C*). This mutual suppression indicates that Ipk2 and Kcs1 act within the same inositol polyphosphate pathway, but argues against a model in which the sensitivity of both mutants reflects simple depletion of a common downstream product; it might suggest instead that residual Kcs1 activity on precursors that accumulate when the pathway is blocked upstream contributes to the *ipk2Δ* phenotype. Expression of wild-type Kcs1-3xFLAG completely restored resistance to zeocin in *kcs1Δ* cells, whereas the kinase-dead Kcs1^K778A^-3xFLAG mutant failed to do so (Fig. 2*D*), demonstrating that Kcs1 catalytic activity, and consequently Kcs1-dependent PP-IPs, are required for resistance to zeocin-induced DNA damage.

**Figure 2.**
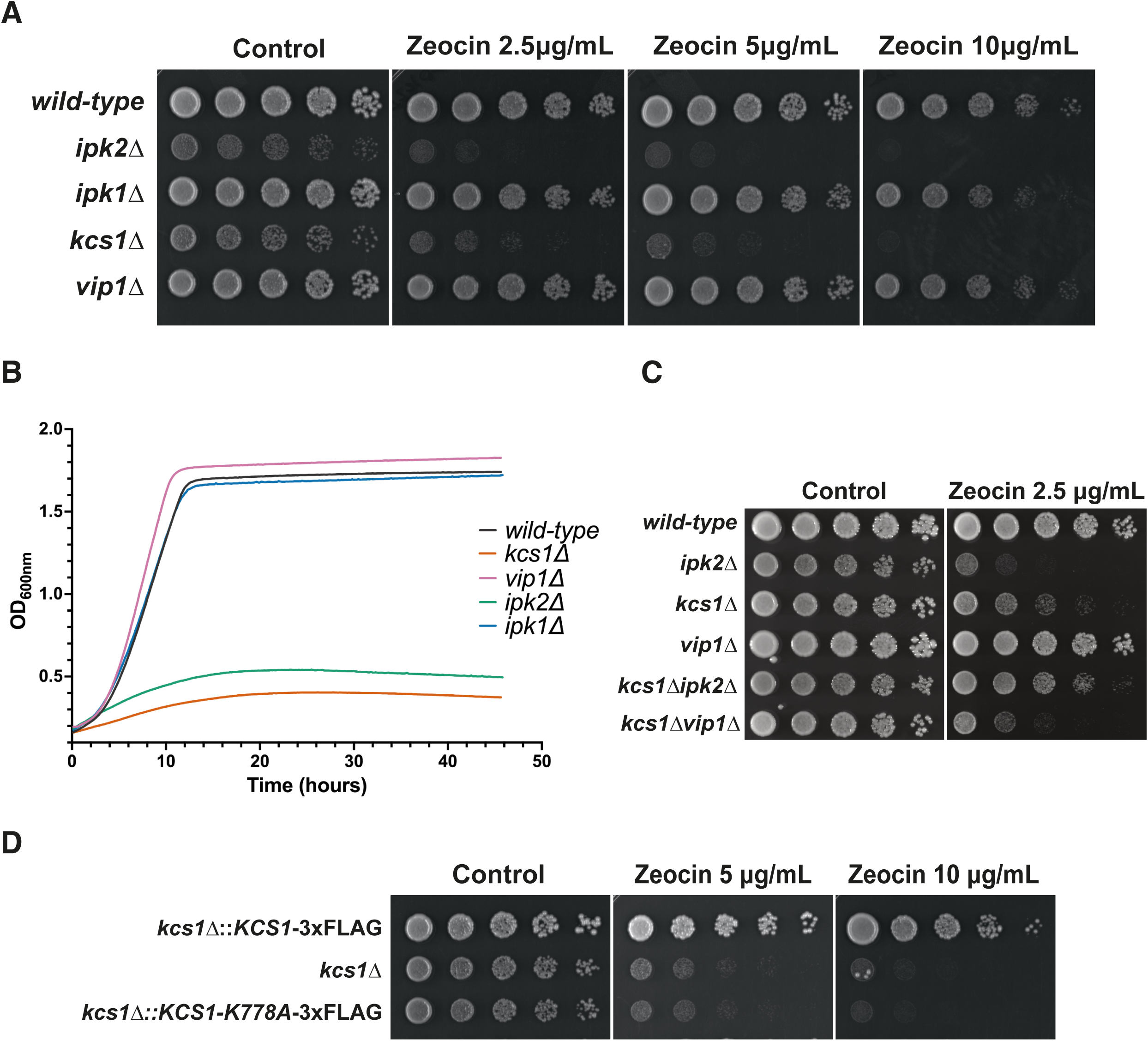
Kcs1-dependent PP-IPs are required for resistance to zeocin-induced genotoxic stress. (A) Five-fold serial dilutions of exponentially growing wild-type, *ipk1Δ*, *ipk2Δ*, *kcs1Δ* and *vip1Δ* cells were spotted onto YPD plates containing the indicated concentrations of zeocin, or onto drug-free YPD (control), and incubated at 30°C for 2–3 days. (B) Growth curves of the indicated yeast strains were generated by monitoring OD600 over time (YPD medium + zeocin 10 μg/mL), as described in the Experimental procedures section. Each line represents the median of two biological replicates. (C) Five-fold serial dilutions of wild-type, *ipk2Δ*, *kcs1Δ*, *vip1Δ*, *kcs1Δipk2Δ* and *kcs1Δvip1Δ*, spotted and incubated as in (A). (D) Five-fold serial dilutions of *kcs1Δ* and *kcs1Δ* strains expressing wild-type Kcs1-3xFLAG or kinase-dead Kcs1^K778A^-3xFLAG were spotted onto YPD plates containing zeocin. Plates were incubated at 30°C for 2–3 days.

### Kcs1-dependent PP-IPs facilitate DNA damage signaling downregulation and DSB repair

We next asked whether zeocin hypersensitivity in *kcs1Δ* cells results from defective DNA damage checkpoint signaling rather than impaired DNA repair. Analysis of Rad53 phosphorylation revealed earlier and stronger checkpoint activation in *kcs1Δ* cells, which was accompanied by elevated γ-H2A levels compared with wild-type cells (Fig. 3*A*). Because the hypomorphic *rad53-R605A* allele suppresses MMS sensitivity in strains lacking negative regulators of Rad53 (47), we introduced this allele into the *kcs1Δ* background. Although the *kcs1Δrad53-R605A* mutant displayed reduced Rad53 activation, it remained hypersensitive to zeocin and retained high γ-H2A levels (Fig. 3*B* and *C*). Similarly, deletion of *TEL1*, the yeast ortholog of mammalian ATM involved in Rad53 activation following DSB formation, also failed to rescue zeocin sensitivity (Fig. S2). These findings indicate that persistent Rad53 hyperactivation in *kcs1Δ* cells is more likely a consequence of defective DNA repair than its primary cause.

**Figure 3.**
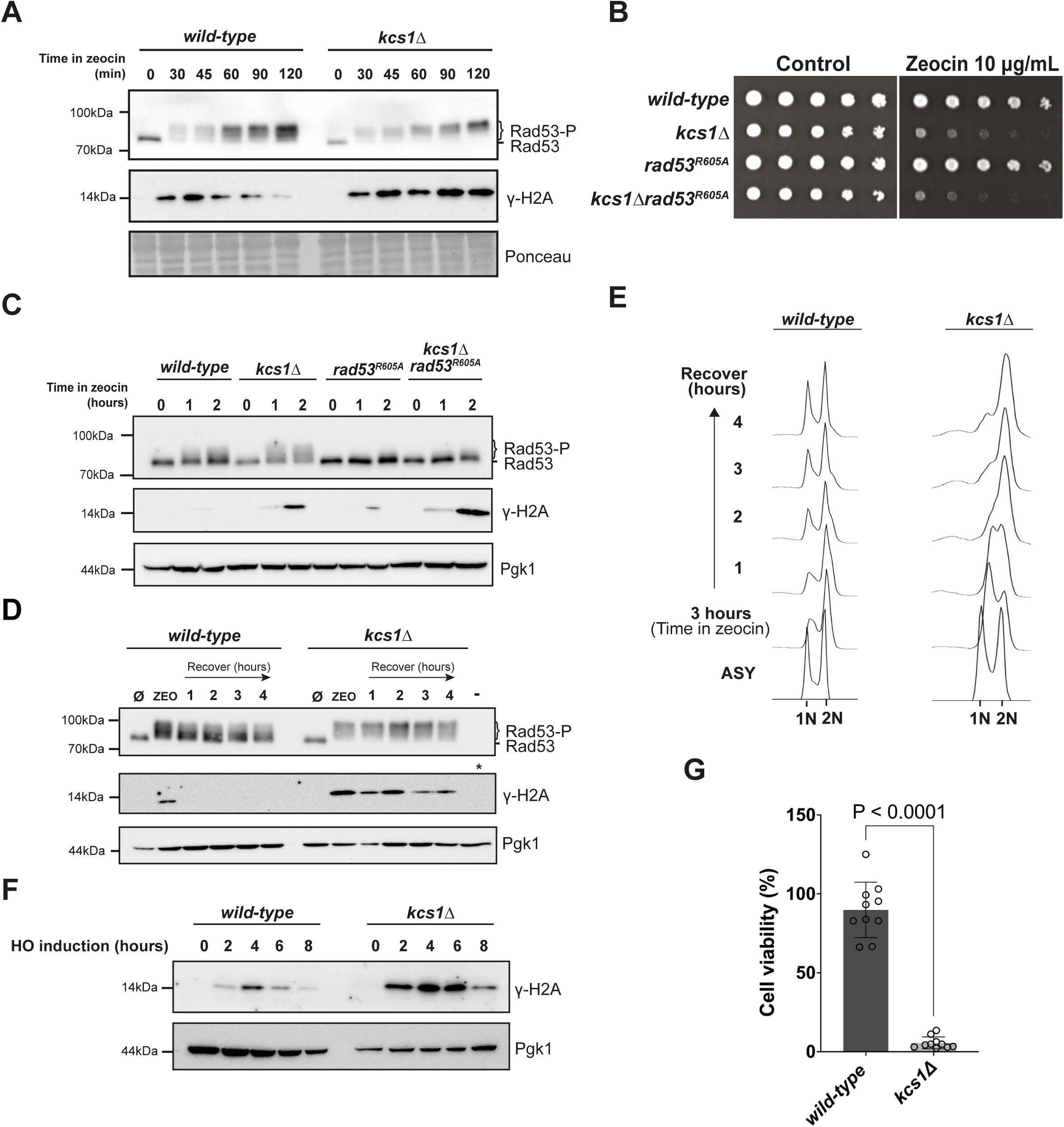
Loss of Kcs1 causes persistent DNA damage signaling and impaired recovery after DSB induction. (A) Exponentially growing wild-type and *kcs1Δ* cells were treated with zeocin (100 µg/mL) and collected at the indicated times. Rad53 phosphorylation (Rad53-P) and γ-H2A were analyzed by western blotting. Molecular mass markers are indicated on the left. (B) Five-fold serial dilutions of exponentially growing wild-type, *kcs1Δ*, *rad53-R605A* and *kcs1Δrad53-R605A* cells were spotted onto YPD plates containing the indicated concentrations of zeocin, or onto drug-free YPD (control), and incubated at 30°C for 2–3 days. (C) The strains in (B) were treated with zeocin and analyzed as in (A). (D) Scheme of the zeocin treatment-and-recovery protocol used in (E) and (F). (E) wild-type and *kcs1Δ* cells were treated with zeocin (100 µg/mL) for 3 h, washed, and released into drug-free YPD. Samples collected before treatment (0), at the end of treatment, and 1, 2, 3 and 4 h after release were analyzed as in (A). (F) DNA content of the samples in (E) was determined by flow cytometry; 1N and 2N positions are indicated. (G) A site-specific DSB was induced by galactose-dependent expression of HO endonuclease in the YRA213 reporter strain, and γ-H2A levels were analyzed at the indicated time points. (H) Survival after HO induction, calculated as the ratio of colonies formed on YPGal to colonies formed on YPD after 72 h at 30°C. Bars represent the mean ± SEM of 12 independent biological replicates per strain; symbols show individual replicates.

Recovery experiments revealed persistent DDR signaling in the absence of Kcs1. Following zeocin removal, *kcs1Δ* cells maintained elevated Rad53 phosphorylation and γ-H2A levels for up to four hours, whereas wild-type cells efficiently attenuated DDR signaling (Fig. 3*D*). Flow cytometry further showed that wild-type cells, as expected, accumulated in G2/M during zeocin treatment and resumed cell cycle progression upon recovery, whereas *kcs1Δ* cells remained arrested in G1/S phase before accumulating in G2/M after zeocin removal, consistent with unresolved DNA damage leading to persistent checkpoint activation (Fig. 3*E*). Similar defects were observed in *ipk2Δ* cells, which also exhibited sustained Rad53 activation and elevated γ-H2A levels during recovery (Fig. S3), supporting the idea that the repair defect results from depletion of PP-IPs rather than their precursors (IP4, IP5 or IP6).

Zeocin belongs to the bleomycin/phleomycin family of radiomimetic antibiotics, which damage DNA through metal-dependent, free-radical attack on the deoxyribose backbone, generating a heterogeneous mixture of single-strand breaks, abasic sites, and double-strand breaks with chemically blocked, non-ligatable termini, of which only a minor fraction (∼10%) are DSB-forming lesions (48, 49). Because of this heterogeneity, and because zeocin-induced damage also has a diffusible, ROS-mediated component (49), phenotypes observed after zeocin treatment cannot be unambiguously attributed to a defect in DSB repair per se. To directly test this, we used a strain expressing the site-specific HO endonuclease under the control of the *GAL1* promoter, which generates a single, chemically defined DSB with termini immediately compatible with both HR and NHEJ, and asked whether the hypersensitivity of *kcs1Δ* cells specifically reflects defective DSB repair. *KCS1* deletion resulted in marked γ-H2A accumulation following DSB induction (Fig. 3*F*), which was accompanied by a substantial loss of cell viability (Fig. 3*G*). Because the HO system delivers a single, chemically defined DSB independently of drug uptake, metal chemistry or cell-wall permeability, these phenotypes cannot be attributed to increased lesion formation in *kcs1Δ* cells. These findings indicate that Kcs1-dependent PP-IPs are required for efficient DSB repair and proper termination of DDR signaling.

### Kcs1 promotes DSB repair independently of HR

Since zeocin-induced DSBs are primarily repaired by HR (50), and Kcs1 has previously been implicated in efficient HR-mediated DSB repair in both yeast and mammals (29, 30), we examined the genetic interaction between Kcs1 and core members of the *RAD52* epistasis group. Both *kcs1Δrad51Δ* and *kcs1Δrad54Δ* double mutants displayed additive sensitivity to zeocin (Fig. 4*A* and *B*), indicating that Kcs1 promotes genome stability independently of core HR proteins. To assess HR directly, we monitored Rad52-mRuby2 focus formation after zeocin treatment. In asynchronous cultures, *kcs1Δ* cells showed a marked reduction in the number of Rad52 foci relative to wild-type after zeocin treatment for up to 4 hours (Fig. S4), an observation that at first sight matched the HR defect previously reported for IP6K1-deficient mammalian cells (29). However, *kcs1Δ* cells arrest in G1/S under zeocin treatment (Fig. 3*E*), whereas HR operates only in S and G2/M. A lower focus count in an asynchronous population could therefore reflect cell-cycle distribution rather than an intrinsic inability to assemble HR intermediates. To separate these possibilities, we repeated the experiment in nocodazole-arrested cells, so that both strains were uniformly in G2/M at the time of damage. In nocodazole-arrested G2/M cells, zeocin treatment induced Rad52 foci in both strains, with *kcs1Δ* cells showing a higher mean proportion of foci-positive cells than wild-type (Fig. 4*C* and *D*). Although this difference did not reach statistical significance, the data show that *kcs1Δ* cells retain the ability to load Rad52 at damage sites, arguing against a defect in the early steps of homologous recombination. The trend toward more foci is consistent with a larger amount of unrepaired lesions in the absence of Kcs1, indicating that recruitment of the HR machinery to damage is not impaired by loss of Kcs1 and that the apparent defect in asynchronous cultures results from altered cell-cycle distribution. In line with this conclusion, synchronized *kcs1Δ* cells (Fig. S5) accumulated higher γ-H2A levels than wild-type cells following zeocin treatment in both G1 and G2/M (Fig. 4*E*), indicating that Kcs1 promotes DSB repair independently of HR, which is inactive during G1 in haploid yeast.

**Figure 4.**
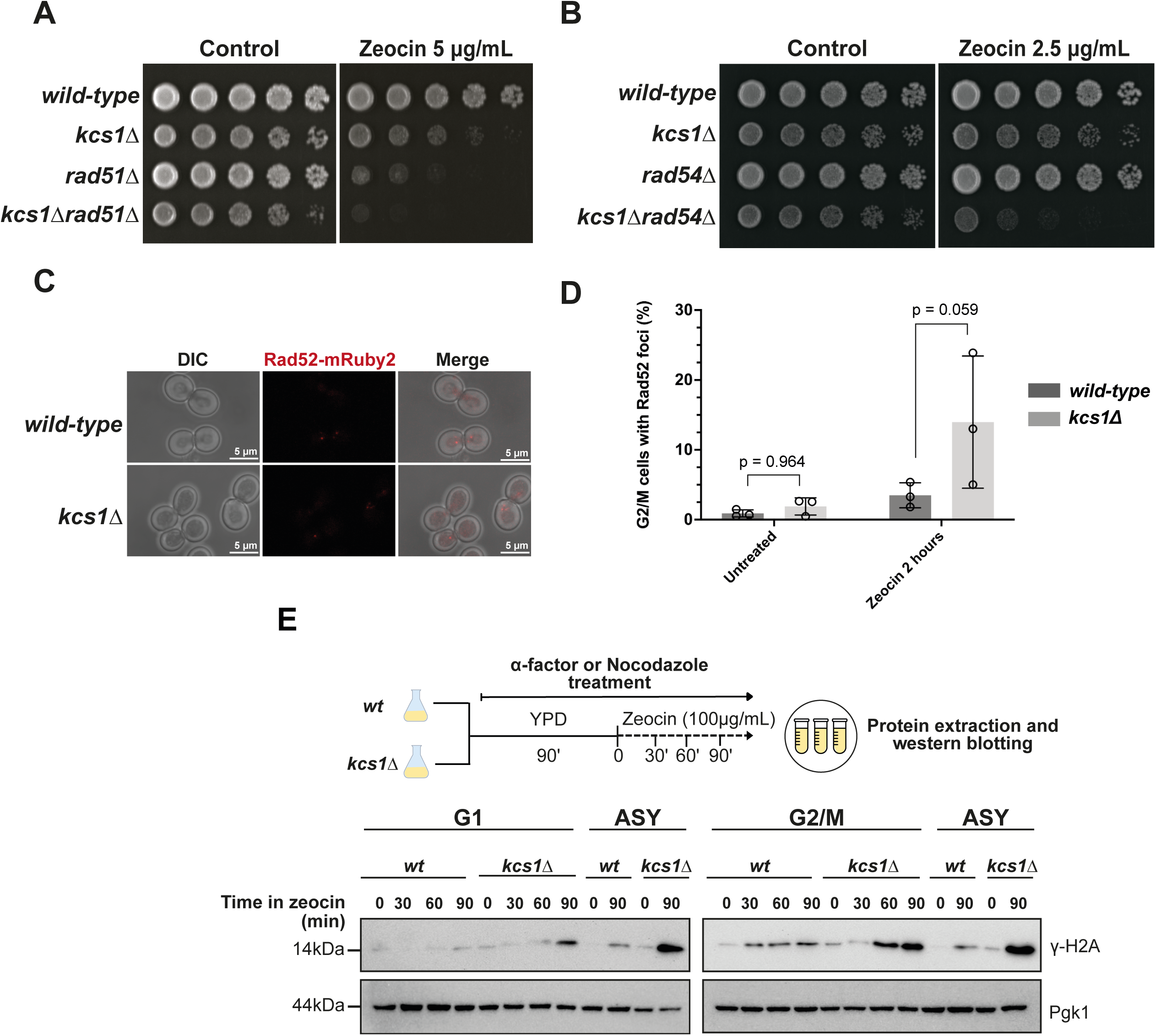
Kcs1 promotes DSB repair independently of the core HR machinery. (A) Five-fold serial dilutions of wild-type, *kcs1Δ*, *rad51Δ*, and *kcs1Δrad51Δ* strains were spotted onto YPD plates containing the indicated concentrations of zeocin. (B) Five-fold serial dilutions of wild-type, *kcs1Δ*, *rad54Δ*, and *kcs1Δrad54Δ* strains were spotted onto YPD plates containing zeocin. (C) Representative images of Rad52-mRuby2 foci in wild-type and *kcs1Δ* cells. (D) Quantification of Rad52-mRuby2 nuclear foci in nocodazole-arrested (G2/M) cells. wild-type and *kcs1Δ* cells were arrested with nocodazole (10 µg/mL, 2 h), then treated with zeocin (100 µg/mL) for 2 h or left untreated and imaged by confocal microscopy. Bars represent the mean percentage of cells containing at least one Rad52-mRuby2 focus from three independent biological replicates (untreated: *wild-type* 0.91 ± 0.51%, *kcs1Δ* 1.89 ± 1.20%; zeocin: *wild-type* 3.49 ± 1.79%, *kcs1Δ* 13.97 ± 9.55%). Individual replicate values are shown as symbols. At least 150 cells were scored per strain and condition in each replicate. Statistical analysis was performed by two-way ANOVA matched on both factors (genotype and treatment), with Sidak’s correction for multiple comparisons applied to genotype comparisons within each treatment and using separate variances. Quantitative data are presented as mean ± SD from the indicated number of independent experiments. (E) γ-H2A levels were analyzed by western blotting following zeocin treatment of cells arrested in G1 or G2/M.

### Kcs1 promotes DSB repair through activation of the Pho80-Pho85 CDK complex

To identify factors downstream of Kcs1 that contribute to DSB repair, we performed SILAC-based quantitative proteomics. Comparison of wild-type and *kcs1Δ* cells following zeocin treatment identified numerous proteins with altered abundance (Fig. 5*A* and File S1). The experiment comprised two independent biological replicates with reciprocal (forward and reverse) SILAC labeling, and only proteins showing reproducible changes in both labeling directions were retained. Kcs1 itself was the most strongly depleted protein in *kcs1Δ* cells (Fig. 5*A*), providing an internal control for the quantification. No reproducible change was detected for the core homologous recombination and checkpoint machinery, although a small number of DNA metabolic enzymes, including the apurinic/apyrimidinic endonuclease Apn1 and topoisomerase I, were reduced in *kcs1Δ* cells (File S1).

**Figure 5.**
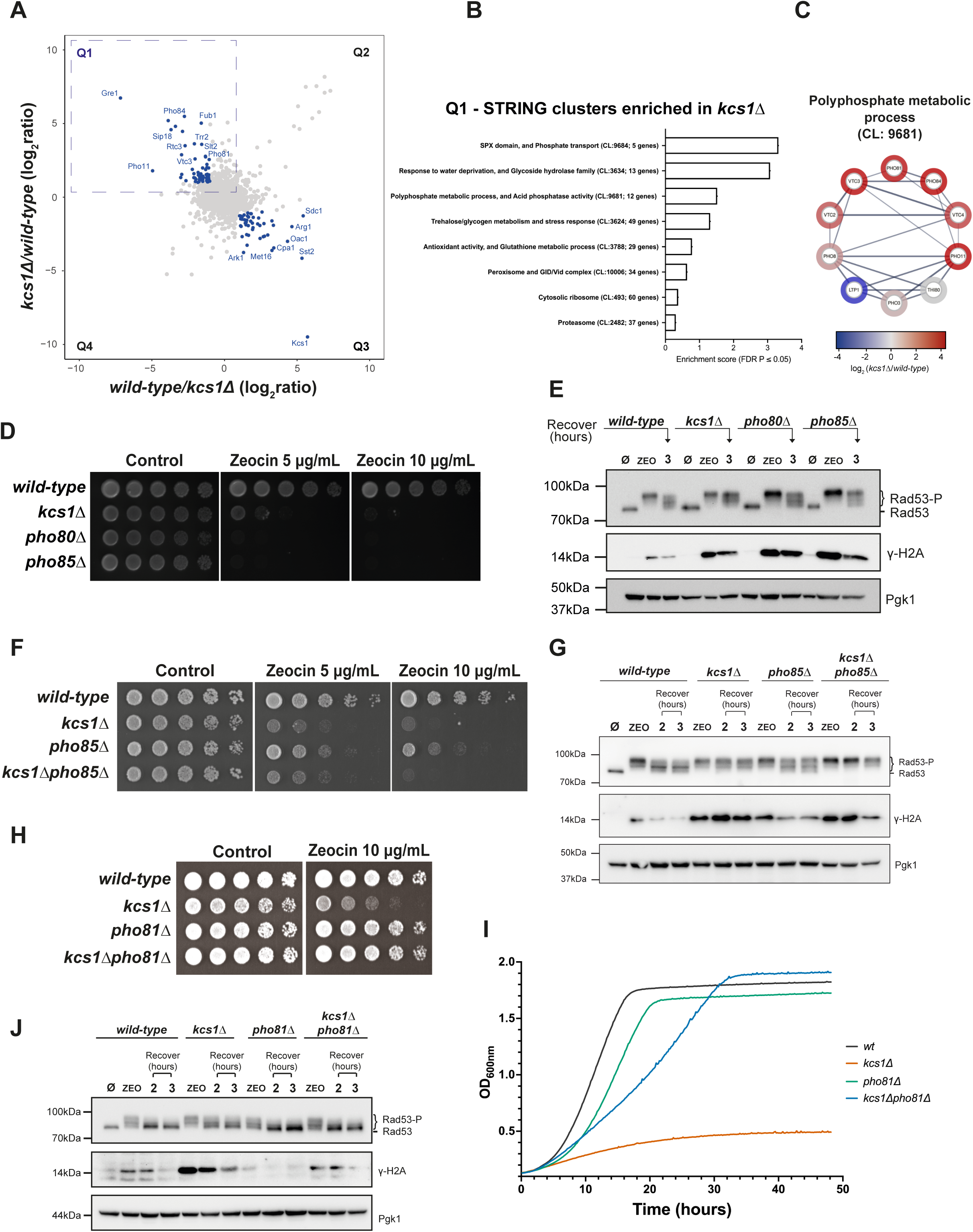
The Pho80-Pho85 cyclin-dependent kinase functions downstream of Kcs1 in the DNA damage response. (A) SILAC-based quantitative proteomic analysis of wild-type and *kcs1Δ* cells treated with zeocin. Proteins showing altered abundance in *kcs1Δ* cells are indicated. (B) Functional modules enriched among proteins increased in *kcs1Δ* cells. Rank-based enrichment of all 2,674 quantified proteins against STRING local network clusters. STRING cluster identifiers and the number of member proteins quantified here are given in parentheses. Two pairs of bars describe the same module at two levels of hierarchy: CL:9684 is the SPX-domain core of CL:9681, and CL:3634 is nested within CL:3624. All clusters shown pass FDR ≤ 1.8 × 10⁻³. (C) STRING association network of the 12 proteins of local network cluster CL:9681 (Polyphosphate metabolic process, and Acid phosphatase activity), one of the clusters enriched in panel B. Edges represent STRING associations at medium confidence (combined score ≥ 0.400) and edge thickness is proportional to the confidence score. Disconnected nodes are not depicted in the figure. Node color encodes the mean log2 ratio measured in this experiment (color scale) with red denoting increased and blue decreased abundance in *kcs1Δ*. (D) Five-fold serial dilutions of wild-type, *pho80Δ*, and *pho85Δ* strains were spotted onto YPD plates containing the indicated concentrations of zeocin. (E) Rad53 phosphorylation and γ-H2A levels were analyzed by western blotting in wild-type, *pho80Δ*, and *pho85Δ* cells following zeocin treatment. (F) Five-fold serial dilutions of *kcs1Δ, pho85Δ,* and *kcs1Δpho85Δ* strains were spotted onto YPD plates containing zeocin. (G) Rad53 phosphorylation and γ-H2A levels were analyzed by western blotting in the indicated strains following zeocin treatment. (H) Five-fold serial dilutions of wild-type, *kcs1Δ, pho81Δ,* and *kcs1Δpho81Δ* strains were spotted onto YPD plates containing zeocin. (I) Growth curves of the indicated yeast strains were generated by monitoring OD600 over time (YPD medium + zeocin 10 μg/mL), as described in the Experimental procedures section. Each line represents the median of two biological replicates. (J) Rad53 phosphorylation and γ-H2A levels were analyzed by western blotting in the indicated strains following zeocin treatment. For (D), (F) and (H), plates were incubated at 30°C for 2–3 days.

To identify the cellular processes affected, we ranked all 2,674 quantified proteins by their mean log2 ratio and tested for functional enrichment against STRING local network clusters (https://string-db.org), an approach that uses the entire quantified proteome and requires no abundance cut-off (Fig. 5*B*). Proteins increased in *kcs1Δ* were enriched for clusters comprising the trehalose/glycogen and general stress response, antioxidant and glutathione metabolism, the proteasome and the cytosolic ribosome, and for phosphate transport and polyphosphate metabolism. Conversely, proteins decreased in *kcs1Δ* were enriched for amino acid biosynthesis, including the methionine and sulfur assimilation, arginine, leucine, lysine and histidine pathways, a composition characteristic of the general amino acid control (Gcn4) regulon, as well as for nucleotide and folate metabolism, RNA polymerase II transcription initiation and P-body components (Fig. S6). The phosphate signature was the most prominent feature of the dataset. The SPX cluster comprises Pho81, the high-affinity phosphate transporter Pho84 and three subunits of the vacuolar transporter chaperone (VTC) polyphosphate polymerase complex, all increased in *kcs1Δ* (Fig. 5*C*), with Pho84, Pho11, Vtc3 and Pho81 among the twenty-one most increased proteins of the 2,674 quantified (File S1). Because Pho80-Pho85 represses the PHO regulon by phosphorylating Pho4 and driving it out of the nucleus, increased abundance of PHO pathway proteins is the expected transcriptional output of a kinase complex that is constitutively inhibited (34, 35). This is consistent with the well-established role of PP-IPs in regulating phosphate homeostasis through Pho80-Pho85 (3, 18). These findings therefore indicate that Pho80-Pho85 is constitutively inhibited in *kcs1Δ* cells and raise the possibility that this defect contributes to impaired DSB repair.

In line with this hypothesis, deletion of either *PHO80* or *PHO85* recapitulated the zeocin sensitivity and DDR defects observed in *kcs1Δ* cells (Fig. 5*D* and *E*). Because Pho85 associates with multiple cyclins controlling diverse cellular processes (40), we next tested whether this phenotype was specific to Pho80. Deletion of other *PHO85* cyclin partners failed to confer zeocin sensitivity, supporting the conclusion that Pho80 is the relevant cyclin required for resistance to DSB-inducing agents (Fig. S7). To determine whether Kcs1 and Pho85 function in the same pathway, we generated the *kcs1Δpho85Δ* double mutant. Deletion of *PHO85* did not further increase zeocin sensitivity in *kcs1Δ* cells (Fig. 5*F*), and the double mutant displayed Rad53 phosphorylation and γ-H2A levels comparable to those of either single mutant (Fig. 5*G*). These results support a model in which Kcs1-dependent PP-IPs promote DSB repair through Pho80-Pho85 signaling.

To test whether inhibition of Pho80-Pho85 activity underlies the DSB repair defect in *kcs1Δ* cells, we deleted the CDK inhibitor *PHO81* in the *kcs1Δ* background with the expectation that it would alleviate the Pho80-Pho85 inhibition, leading to resistance to genotoxic stress induced by zeocin. In line with that, deletion of *PHO81* significantly rescued zeocin sensitivity and restored normal γ-H2A levels and Rad53 activation kinetics in *kcs1Δ* cells (Fig. 5*H*-*J*), indicating that constitutive inhibition of the Pho80-Pho85 kinase complex is responsible for the DNA repair defects associated with PP-IP depletion. Together, these findings support a model in which Kcs1-dependent PP-IPs promote DSB repair by maintaining Pho80-Pho85 activity.

### Kcs1 promotes DSB repair independently of phosphate homeostasis

Pho80 is the cyclin that partners with Pho85 to regulate the PHO pathway in response to phosphate availability (34, 35). Therefore, we asked whether altered phosphate metabolism could account for the impaired DSB repair observed in *kcs1Δ* cells. It has been shown that deletion of *KCS1* causes an increase in the concentration of inorganic phosphate (Pi) both in yeast and mammals (19, 39). If constitutive activation of the PHO pathway were responsible, reducing phosphate signaling or uptake would be expected to rescue the phenotype. However, although deletion of the transcription factor Pho4 partially restored zeocin resistance, deletion of PHO pathway targets, including *PHO84* and *PHO5*, failed to rescue the sensitivity of *kcs1Δ* cells (Fig. 6*A*). Likewise, phosphate depletion from the growth medium did not restore zeocin resistance (Fig. 6*B*). Because Kcs1 is also required for polyphosphate accumulation in acidocalcisomes (51, 52), we next asked whether polyphosphate depletion contributes to the repair defect. However, unlike *kcs1Δ* cells, a *vtc4Δ* strain, which is defective in polyphosphate synthesis, showed no increase in zeocin sensitivity (Fig. 6*C*). Together, these findings demonstrate that the role of Kcs1-dependent PP-IPs in DSB repair cannot be explained by alterations in phosphate homeostasis, pointing instead to a previously unrecognized function of the Pho80-Pho85 signaling axis during the DNA damage response.

**Figure 6.**
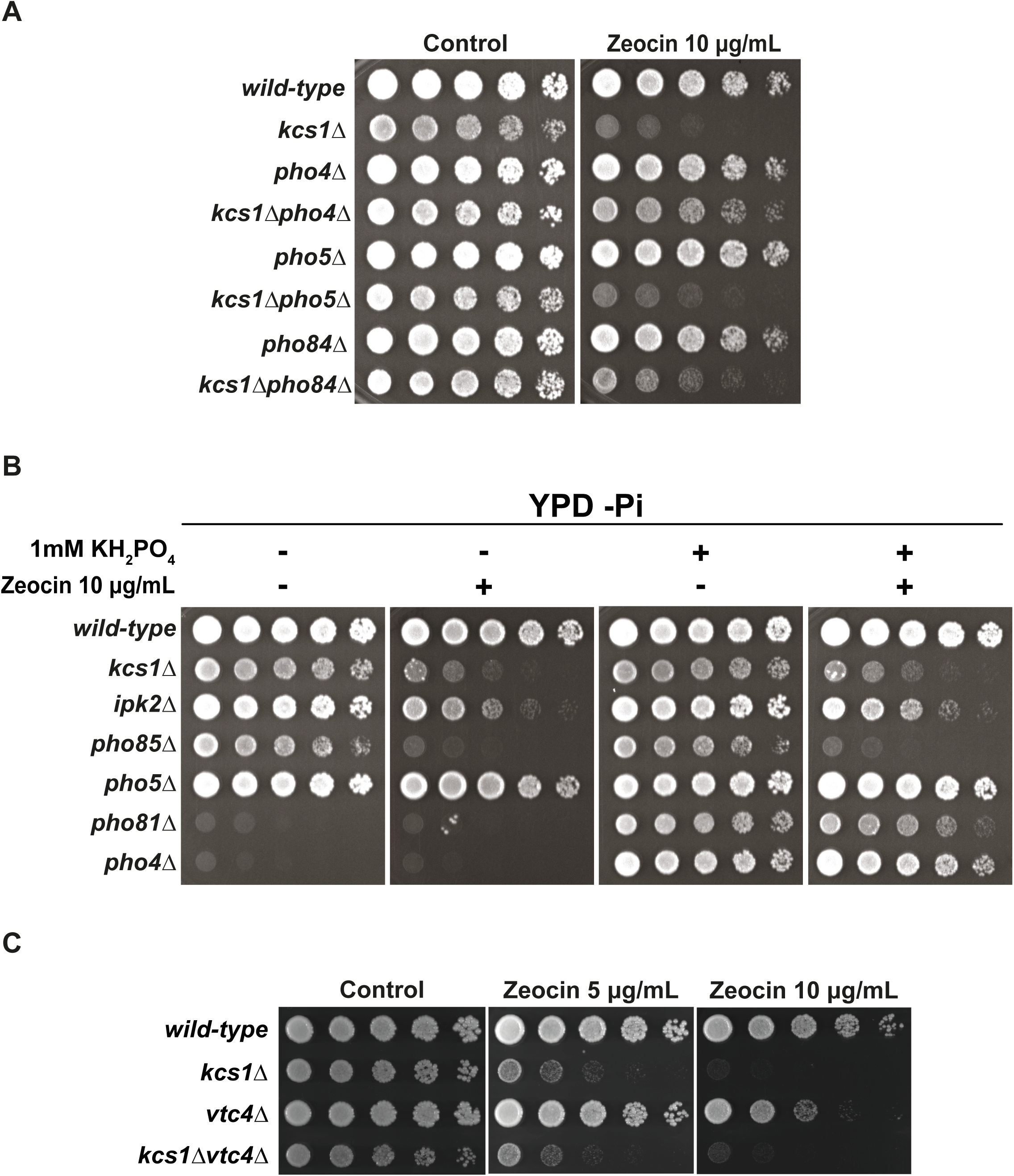
The Kcs1-dependent DSB repair defect is independent of phosphate homeostasis. (A) Five-fold serial dilutions of wild-type, *kcs1Δ*, and the indicated kcs1Δ strains carrying deletions of genes involved in phosphate sensing or acquisition were spotted onto YPD plates containing zeocin. (B) wild-type and mutant strains were grown under the indicated phosphate concentrations, and sensitivity to zeocin was assessed by serial dilution. (C) Five-fold serial dilutions of wild-type, *kcs1Δ*, *vtc4Δ* and *kcs1Δvtc4Δ* strains were spotted onto YPD plates containing the indicated concentrations of zeocin. Plates were incubated at 30°C for 2–3 days.

### Ku80 deletion suppresses the DNA repair defects of kcs1Δ cells

Because alterations in phosphate homeostasis do not explain the DSB repair defects observed in *kcs1Δ* cells, we next investigated other potential Pho85 targets. Pho85 has been proposed to phosphorylate Ku80 through its association with the cyclin Pcl1 during S phase and G2/M, thereby limiting NHEJ and favoring HR (53). Intriguingly, deletion of *YKU80*, but not *YKU70*, substantially rescued the zeocin sensitivity of *kcs1Δ* cells (Fig. 7*A* and *B*). Moreover, *YKU80* deletion restored checkpoint deactivation and reduced γ-H2A accumulation (Fig. 7*C*). To determine whether this phenotype was dependent on Pho85, we deleted *YKU80* in a *pho85Δ* background. In contrast to *kcs1Δ* cells, deletion of *YKU80* did not rescue the zeocin sensitivity of *pho85Δ* cells (Fig. 7*D*). Nuclear actin has recently been shown to drive INO80-dependent conversion of zeocin-induced lesions into DSBs in budding yeast (54, 55) and F-actin influences Ku70-Ku80 retention at DNA lesions and Ku-dependent NHEJ efficiency in mammalian cells (56–58). We therefore asked whether Kcs1-dependent PP-IPs act on DSB repair through actin dynamics. Treatment with the actin-depolymerizing drug Latrunculin A (LatA), which raises nuclear actin levels, did not suppress but instead further increased γ-H2A levels in zeocin-treated *kcs1Δ* cells (Fig. 7*E*), indicating that the two inputs act additively rather than within the same pathway. LatA alone was not included, as actin depolymerization does not by itself induce γ-H2A accumulation or checkpoint activation in the absence of a damaging agent (55).

**Figure 7.**
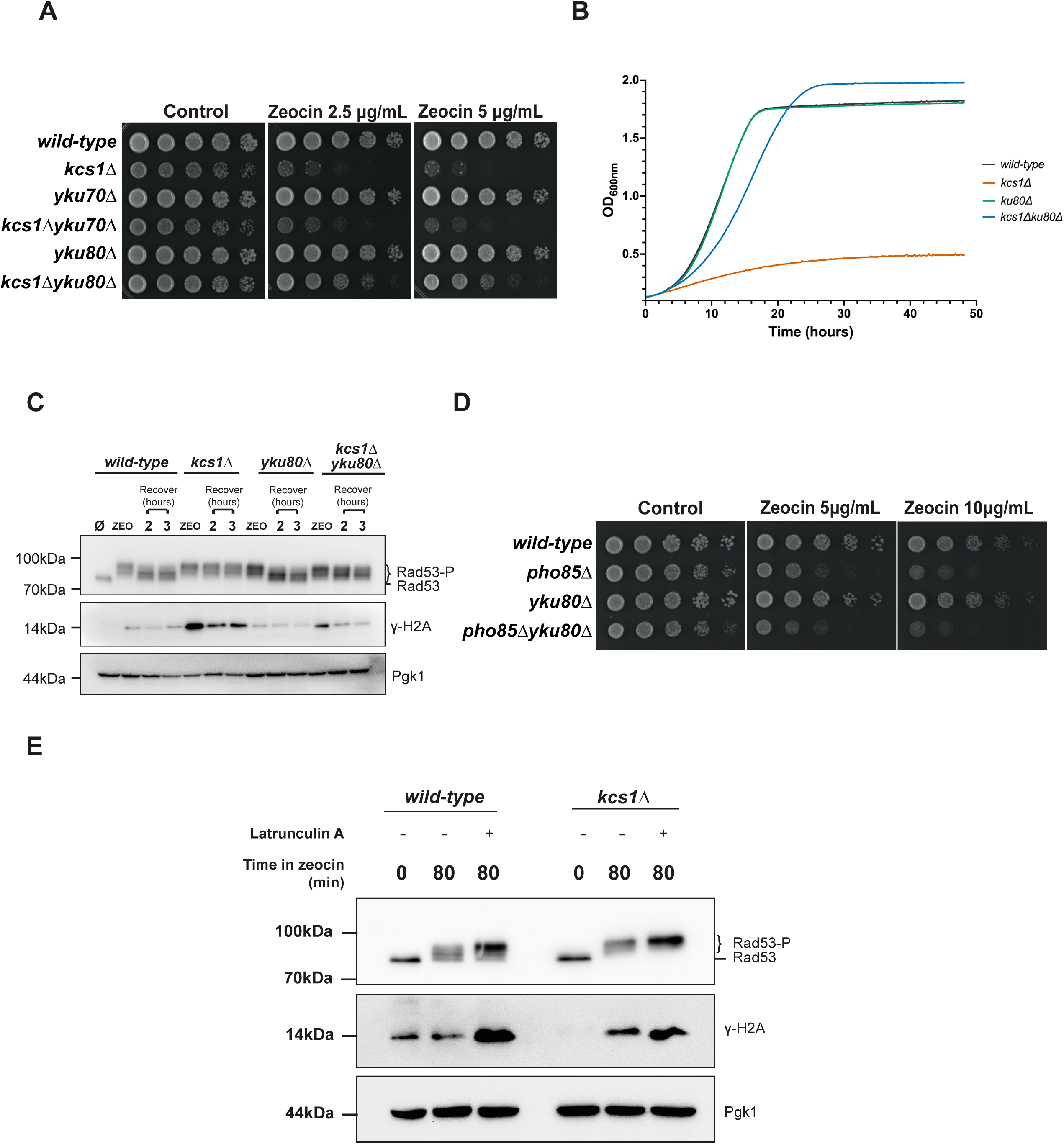
Deletion of *YKU80* suppresses the DNA damage sensitivity of *kcs1Δ* cells. (A) Five-fold serial dilutions of wild-type, *kcs1Δ, yku70Δ, yku80Δ, kcs1Δyku80Δ* and *kcs1Δyku70Δ* strains were spotted onto YPD plates containing the indicated concentrations of zeocin. (B) Growth curves of the indicated yeast strains in the presence of zeocin 10 μg/mL were generated by monitoring OD600 over time, as described in the Experimental procedures section. Each line represents the median of two biological replicates. (C) Rad53 phosphorylation and γ-H2A levels were analyzed by western blotting in wild-type, *kcs1Δ, yku80Δ,* and *kcs1Δyku80Δ* cells following zeocin treatment. (D) Five-fold serial dilutions of the indicated *pho85Δ, yku80Δ*, and double-mutant strains were spotted onto YPD plates containing zeocin. Plates were incubated at 30°C for 2–3 days. (E) wild-type and *kcs1Δ* cells were treated with zeocin (50 µg/mL) alone or in combination with Latrunculin A (20 µM) for 80 min, and Rad53 phosphorylation and γ-H2A levels were analyzed by western blotting.

Thus, although loss of Ku80 suppresses the DNA damage phenotypes associated with loss of Kcs1, this effect cannot be explained solely by the proposed Pho85-dependent phosphorylation of Ku80. These findings raise the possibility that Kcs1-dependent PP-IPs influence Ku80 function through an NHEJ-independent mechanism or through additional Pho85-independent processes.

## Discussion

In this study, we identify Kcs1-dependent PP-IPs as critical regulators of the response to DSBs in budding yeast. Loss of Kcs1 causes hypersensitivity to the radiomimetic drug zeocin and to HO endonuclease-induced DSBs, together with persistent checkpoint activation, defective cell-cycle recovery, and elevated γ-H2A levels. These defects reflect a role for Kcs1-dependent PP-IPs in DSB repair that is independent of cell-cycle stage and of the core HR machinery. Using quantitative proteomics, we identify the Pho80-Pho85 cyclin-dependent kinase complex as a major downstream effector of this pathway. In addition, an unexpected genetic interaction between *KCS1* and *YKU80* implicates Ku80-dependent chromatin functions downstream of PP-IP signaling, and this phenotype is further increased, in an additive rather than epistatic manner, by pharmacological disruption of the actin cytoskeleton, pointing to multiple, genetically separable layers of Ku80 regulation.

### Kcs1-dependent PP-IPs are required for DSB repair

In budding yeast, Kcs1 and Vip1 generate distinct inositol pyrophosphate species. Kcs1 preferentially synthesizes 5-IP7, whereas Vip1 produces 1-IP7, and the concerted activity of both enzymes is required for 1,5-IP8 synthesis (3, 4, 59). Deletion of *KCS1* depletes both 5-IP7 and 1,5-IP8 without affecting 1-IP7 levels (37). Our genetic data indicate that the PP-IP species involved in the DDR is generated by Kcs1, as deletion of *KCS1*, but not *VIP1*, caused hypersensitivity to zeocin (Fig. 2*A* and *B*), and this phenotype required Kcs1 catalytic activity (Fig. 2*D*). However, this interpretation is complicated by the observation that *ipk1Δ* cells do not display zeocin sensitivity (Fig. 2*A* and *B*), despite lacking IP6, the precursor required for 5-IP7 synthesis. One possible explanation is that 5-IP7 remains the principal signaling molecule regulating the DDR, whereas elevated 5-PP-IP4 levels in *ipk1Δ* cells compensate for the loss of 5-IP7, since this PP-IP accumulates in the absence of Ipk1 (23). Alternatively, 5-PP-IP4 itself may function as the relevant signaling molecule controlling the DDR. Notably, the absence of zeocin sensitivity in *ipk1Δ* cells also argues against a direct role for IP6 itself in this pathway, despite the established IP6–Ku70/80 interaction described in mammalian cells (28, 43), and points instead to a pyrophosphorylated species as the relevant effector. Distinguishing between these possibilities will require direct quantification of PP-IP species under DNA damage conditions and genetic or biochemical approaches capable of selectively manipulating individual PP-IP pools.

The similar phenotypes shared by *kcs1Δ* and *ipk2Δ* with respect to zeocin sensitivity (Fig. 2*A* and *B*) and increased Rad53 activation and γ-H2A levels (Fig. 3*D* and Fig. S3) further support the conclusion that these enzymes function in the same biosynthetic pathway to generate the Kcs1-dependent PP-IPs required for efficient DSB repair. This conclusion is consistent with previous studies showing that *kcs1Δ* and *ipk2Δ* cells share additional phenotypes, including increased telomere length and defects in vesicular trafficking, which are absent in *ipk1Δ* mutants (21–23). Accordingly, the DNA repair defects observed in *ipk2Δ* cells are most likely a consequence of Kcs1-dependent PP-IP depletion rather than reduced levels of soluble inositol polyphosphates (IP4, IP5 or IP6) or phosphoinositide signaling. This distinction is particularly relevant because recent studies have implicated nuclear phosphoinositide signaling in the DNA damage response (60). The mammalian Ipk2 ortholog IPMK, for example, promotes ATR activation by generating nuclear PI(3,4,5)P3, thereby stimulating nuclear actin assembly and ATR recruitment to DNA lesions (44). Whether PP-IPs also contribute to this signaling axis has not been addressed. Because PP-IPs can compete with phosphoinositides such as PI(3,4,5)P3 for binding to PH-domain-containing proteins (16), it is conceivable that these signaling molecules cooperate or compete to coordinate chromatin responses during the DDR. However, cells lacking *IPK2*, but not *KCS1*, show sensitivity to H2O2, (Fig. S1), implying that Ipk2 has Kcs1-independent roles that are important for stress resistance. At least two such activities have been described. First, Ipk2/Arg82 possesses nuclear phosphoinositide 3-kinase activity, converting PtdIns(4,5)P2 into PtdIns(3,4,5)P3, and this lipid kinase activity contributes to transcriptional regulation (61). Second, Ipk2/Arg82 acts as a structural chaperone of the MADS-box proteins Mcm1 and Arg80 within the ArgR–Mcm1 transcriptional complex, a function that depends on its polyaspartate domain rather than on its catalytic activity (62, 63). Loss of Ipk2 therefore compromises transcriptional programs that remain intact in *kcs1Δ* cells.

Two features of our genetic analysis deserve emphasis. First, *VIP1* is not fully epistatic with *KCS1*. Although *vip1Δ* cells are not zeocin-sensitive on their own, the *kcs1Δvip1Δ* double mutant is consistently slightly more sensitive than *kcs1Δ* alone (Fig. 2*C*). Vip1 therefore makes a minor but detectable contribution to zeocin resistance that is revealed only once the Kcs1-dependent species are absent. Second, deletion of *IPK2* does not simply phenocopy deletion of *KCS1*: the two mutations suppress each other, so that *kcs1Δipk2Δ* cells are more zeocin-resistant than either single mutant (Fig. 2*C*), even though the double mutant lacks Kcs1-dependent PP-IPs altogether. A possible interpretation is that it is the imbalance of the inositol phosphate pool, rather than the absolute level of any single species, that compromises the response to DSBs: in *kcs1Δ* cells IP6 accumulates because it is no longer converted into 5-IP7, whereas in *ipk2Δ* cells Kcs1 remains active on the precursors that accumulate upstream of the pathway and removing both enzymes eliminates both abnormal pools. Distinguishing depletion from imbalance will ultimately require direct quantification of individual inositol phosphate and pyrophosphate species in these mutants under genotoxic stress.

### Kcs1 promotes DSB repair through an HR-independent mechanism

Previous studies implicated PP-IPs in HR-mediated DNA repair in both yeast and mammalian cells (29, 30). Jadav and colleagues, for example, reported that IP6K1-deficient mouse embryonic fibroblasts display persistent γ-H2AX, RAD51 and BLM foci and prolonged cell-cycle arrest following replication stress, leading the authors to propose a defect in HR (29). At first glance, our data appeared consistent with this model, as *kcs1Δ* cells exhibited persistent DDR activation together with a marked reduction in Rad52 nuclear foci following zeocin treatment in asynchronous cultures (Fig. S4). However, this apparent deficit was abolished when damage was induced in G2/M-arrested cells (Fig. 4*C* and *D*), indicating that it results from the altered cell-cycle distribution of *kcs1Δ* cells rather than from an intrinsic HR defect. This conclusion is strongly supported by our genetic analyses. Loss of *KCS1* displayed additive, rather than epistatic, interactions with *RAD51* and *RAD54* (Fig. 4*A* and *B*) following exposure to zeocin, demonstrating that Kcs1-dependent PP-IPs function independently of the core HR machinery. Likewise, the persistence of elevated γ-H2A levels in G1-arrested *kcs1Δ* cells (Fig. 4*E*) further argues against an HR-specific defect, as canonical HR is inactive during G1 in haploid yeast cells. Together, these findings indicate that Kcs1-dependent PP-IPs promote DSB repair through a mechanism that is independent of HR. It is therefore conceivable that at least some phenotypes previously interpreted as evidence for defective HR in PP-IP-deficient cells may instead arise indirectly from altered cell-cycle progression.

### PP-IPs promote DSB repair through modulation of the Pho80-Pho85 CDK activity

To identify downstream effectors of Kcs1-dependent PP-IP signaling in the DDR, we used SILAC-based quantitative proteomics to compare wild-type and *kcs1Δ* cells after zeocin treatment. To our knowledge, this is the first proteome-wide, quantitative characterization of the Kcs1-dependent proteome. This analysis identified the PHO pathway among the processes most prominently affected in *kcs1Δ* cells (Fig. 5*B* and *C*), providing the starting point for the genetic dissection of the Pho80-Pho85 axis. One of the major findings of this study is that deletion of *PHO81* suppresses the DNA repair and checkpoint defects caused by loss of Kcs1, placing the Pho80-Pho85 kinase complex downstream of Kcs1-dependent PP-IP signaling. Pho81 is the physiological inhibitor of Pho80-Pho85, and recent studies have established that 1,5-IP8 relieves this inhibition by binding to the SPX domain of Pho81, thereby maintaining Pho80-Pho85 activity under phosphate-replete conditions (37).

An important limitation to this model concerns which PP-IP species relieves Pho81-mediated inhibition of Pho80-Pho85 during the DDR. Chabert et al. showed that 1,5-IP8 synthesis requires the combined activities of Kcs1 and Vip1, and that *vip1Δ* cells retain approximately 25% of wild-type 1,5-IP8 levels (a 75% reduction), in contrast to the essentially complete loss of 1,5-IP8 in *kcs1Δ* cells (37). Because *vip1Δ* cells are not zeocin-sensitive and do not substantially enhance the zeocin sensitivity of *kcs1Δ* cells (Fig. 2*A*-*C*), Vip1-dependent synthesis appears dispensable for Kcs1-dependent DSB repair. However, since *vip1Δ* cells retain a substantial residual 1,5-IP8 pool, these data do not formally exclude a contribution of 1,5-IP8 to Pho81 regulation in this context, and instead more directly support a model in which a Kcs1-dependent, Vip1-independent PP-IP species (5-IP7 and/or 5-PP-IP4) is sufficient to activate Pho80-Pho85 upon genotoxic stress. Differences in genetic background between studies may also contribute to differences in the relevant threshold. Directly quantifying PP-IP species following genotoxic stress will help resolve which species is mechanistically responsible.

Although the molecular basis of this regulation remains incompletely understood, our results extend this model by identifying Pho80-Pho85 as a critical regulator of DSB repair. This function appears to be mechanistically distinct from the canonical role of Pho80-Pho85 in phosphate homeostasis, since neither deletion of PHO pathway target genes (Fig. 6*A*) nor phosphate depletion (Fig. 6*B*) restored the DNA repair defects of *kcs1Δ* cells. Similarly, the partial suppression observed in the *kcs1Δpho4Δ* mutant (Fig. 6*A*) is more likely explained by reduced *PHO81* expression, a direct Pho4 target, than by altered phosphate metabolism itself. An alternative explanation for the DNA repair defects observed in *kcs1Δ* cells relates to the severe depletion of vacuolar inorganic polyphosphate (polyP), a well-established consequence of *KCS1* deletion (51, 52). PolyP has been implicated in multiple aspects of genome maintenance in yeast, including cell-cycle progression and chromosome stability (64). However, this possibility is not supported by our genetic analyses: cells lacking Vtc4, the catalytic subunit of the vacuolar polyphosphate polymerase complex, failed to recapitulate the zeocin sensitivity observed in *kcs1Δ* cells (Fig. 6*C*) despite their inability to synthesize polyP. Thus, although *KCS1* deletion profoundly alters phosphate metabolism, neither constitutive PHO activation nor polyP depletion appears sufficient to explain the DSB repair defects observed in *kcs1Δ* cells. Kcs1-dependent PP-IPs were also found to be critical for fungal virulence in *Cryptococcus neoformans*, a human-pathogenic fungus of important public health relevance (32, 65). In this organism, 5-IP7 binding to the SPX domain of Pho81 promotes Pho80-Pho85 activity and is required for proper phosphate homeostasis and virulence (31). Our findings likewise implicate the Pho81–Pho80–Pho85 axis as a critical downstream effector of Kcs1-dependent PP-IP signaling. However, the physiological output of this axis appears to differ between the two fungal systems. Whereas Kcs1-dependent PP-IPs primarily regulate phosphate homeostasis and virulence in *C. neoformans*, our results indicate that their role in DSB repair in budding yeast is not explained by alterations in phosphate metabolism. Determining whether Kcs1-dependent PP-IPs similarly regulate DSB repair in pathogenic fungi will be an important direction for future studies and may reveal conserved mechanisms of genome maintenance that could ultimately be relevant to the development of new antifungal strategies.

### A potential Ku80-centered mechanism to explain DSB repair defects

One of the most unexpected findings of this study was that deletion of *YKU80*, but not *YKU7*0, significantly suppressed the DNA repair defects of *kcs1Δ* cells (Fig. 7*A*). Because canonical NHEJ requires the Ku70-Ku80 heterodimer, this asymmetry argues against a model in which excess or mis-regulated heterodimer activity is responsible for the repair defect, since loss of either subunit should then be equally protective, and points instead to a Ku80-specific function. A previous study proposed that Pcl1-Pho85 phosphorylates Ku80 during S phase and G2/M to suppress inappropriate NHEJ (53). Our genetic data instead suggest that Pho80, rather than Pcl1, is the cyclin relevant to the DNA damage phenotypes associated with loss of Kcs1 (Fig. S7). Notably, deletion of *YKU80* did not rescue the zeocin sensitivity of *pho85Δ* cells (Fig. 7*D*), even though complete loss of Pho85 should eliminate Ku80 phosphorylation more completely than the partial reduction in Pho80-Pho85 activity caused by *KCS1* deletion. This is consistent with the pleiotropic nature of Pho85, which partners with several cyclins beyond Pho80 to control processes such as cell morphogenesis and cytoskeletal organization (40, 66), indicating that complete loss of Pho85 likely compromises multiple processes relevant to genome maintenance, of which Ku80 regulation is only one. However, this interpretation remains tentative because we did not directly examine Ku80 phosphorylation using phospho-deficient mutants. Moreover, the persistence of γ-H2A accumulation in G1-arrested *kcs1Δ* cells (Fig. 4*E*) argues against a simple model in which defective Pho80-dependent Ku80 phosphorylation leads to aberrant NHEJ, since NHEJ is the predominant DSB repair pathway during G1 in haploid yeast. Kcs1-derived PP-IPs may instead, or additionally, regulate Ku80 directly. Structural and biochemical studies demonstrated that IP6 binds to the Ku70-Ku80 heterodimer and stimulates DNA end-joining by promoting assembly and activity of the DNA-PK complex during NHEJ in mammalian cells (28, 42, 43, 67), which raises the possibility that the structurally related PP-IPs modulate Ku80 conformation, protein-protein interactions, or pyrophosphorylation (12). This asymmetry could also reflect a heterodimer-independent pool of Ku80, since subunit loss in mice does not fully eliminate the partner protein, and Ku70 homodimers have been isolated *in vitro* (68, 69). While these mechanisms remain speculative, they offer a plausible framework for the *KCS1*-*YKU80* genetic interaction and warrant future biochemical investigation.

An alternative explanation for the suppression of *kcs1Δ* phenotypes by *YKU80* deletion is that it acts through DNA end resection rather than through a Ku80-specific function. In budding yeast, the Ku70-Ku80 complex antagonizes resection initiation: loss of Ku increases 5′-end degradation at DSBs, an effect that is most pronounced in G1, when CDK activity is low, and largely dispensable in G2, where CDK-driven resection predominates (70). Consistent with this, Ku restricts access of Exo1 and Sgs1-Dna2 to DNA ends, and deletion of *YKU70* restores resection and suppresses the ionizing-radiation and camptothecin sensitivity of resection-defective *sae2Δ* and *mre11* mutants almost to wild-type levels (71). By analogy, if Kcs1-dependent PP-IPs were required for efficient resection initiation, loss of Ku80 could restore end processing through Exo1 and Sgs1 and thereby suppress the sensitivity of *kcs1Δ* cells. Because Ku-mediated end protection is most evident in G1, this model is not excluded by the persistence of γ-H2A in G1-arrested *kcs1Δ* cells. However, a purely resection-based model predicts that deletion of *YKU70* should suppress as efficiently as deletion of *YKU80*, since both subunits are required for Ku to bind DNA ends. The failure of *yku70Δ* to suppress (Fig. 7*A*) therefore argues against this explanation. Discriminating definitively between these possibilities will require direct measurement of resection at an induced DSB, together with epistasis analysis of the suppression with Exo1, Sgs1 and Rad52.

Altered telomere homeostasis is unlikely to explain the *kcs1Δyku80Δ* suppression. Although *yku80Δ* cells have short telomeres (72) and *kcs1Δ* cells long ones (21, 22), *kcs1Δyku70Δ* cells retain the short-telomere phenotype of *yku70Δ* (22) yet remain zeocin-sensitive (Fig. 7*A*). Also, *kcs1Δtel1Δ* cells, in which Kcs1-dependent telomere regulation is abolished (21), are still sensitive (Fig. S2); and extensive telomere elongation per se does not measurably affect yeast fitness or stress resistance (73).

These findings raise the possibility that Ku80 exerts a previously unrecognized, NHEJ-independent function during the DDR that becomes detrimental when Kcs1-dependent PP-IP signaling is disrupted. Notably, neither *PHO81* nor *YKU80* deletion fully restored the zeocin resistance of *kcs1Δ* cells to wild-type levels (Fig. 5*I* and 7*B*), indicating that Kcs1-dependent PP-IPs must regulate additional targets that also contribute to DSB repair. The actin cytoskeleton was an attractive candidate since: (a) F-actin associates with the Ku70-Ku80 complex and is required for its retention at chromatin and for efficient Ku-dependent NHEJ in mammalian cells (57, 58); (b) IP6K-generated PP-IPs regulate actin remodeling through protein binding and pyrophosphorylation (74, 75); and (c) raising nuclear actin with LatA drives INO80-dependent conversion of zeocin-induced lesions into DSBs (54, 55). Had Kcs1-dependent PP-IPs acted through this axis, LatA and *KCS1* deletion should have behaved epistatically. Instead, their effects were additive (Fig. 7*E*), which argues against actin remodeling as the route by which Kcs1-dependent PP-IPs promote DSB repair and places the two inputs in genetically separable pathways. Because Ku proteins have also been implicated in chromatin organization and DNA-end dynamics, it is tempting to speculate that Kcs1-derived PP-IPs coordinate multiple chromatin-associated processes during DSB repair. Defining the molecular targets of Pho80-Pho85 and determining whether PP-IPs directly regulate Ku80 or actin cytoskeletal dynamics will be important goals for future studies.

### A working model for the contribution of the Kcs1/Pho80-Pho85 axis for DSB repair and genomic stability

We propose a model in which Kcs1-dependent PP-IPs function as a key regulator of DSB repair in budding yeast (Fig. 8). Upon genotoxic stress, Kcs1-dependent PP-IPs relieve Pho81-mediated inhibition of Pho80-Pho85, and active Pho80-Pho85 promotes accurate DSB repair through phosphorylation of Ku80, which limits NHEJ in S and G2/M, together with phosphorylation of additional, as yet unidentified, targets that may act through another biological process such as chromatin remodeling, actin cytoskeleton organization and/or another process important for DSB repair. In addition, Kcs1-dependent PP-IPs may act through one or more Pho80-Pho85-independent routes, potentially involving protein pyrophosphorylation or allosteric regulation of protein targets, that contribute additively to DSB repair and would account for the activity of this pathway outside S and G2/M (Fig. 8*A*). In *kcs1Δ* cells, depletion of Kcs1-dependent PP-IPs leaves Pho81 bound to and inhibiting Pho80-Pho85, causing pleiotropic misregulation of Ku80 and other CDK targets and loss of the additive Pho85-independent contribution. Together, these defects lead to accumulation of unresolved DSBs, persistent DNA damage signaling, and G2/M arrest leading to genomic instability and cell death (Fig. 8*B*).

**Figure 8.**
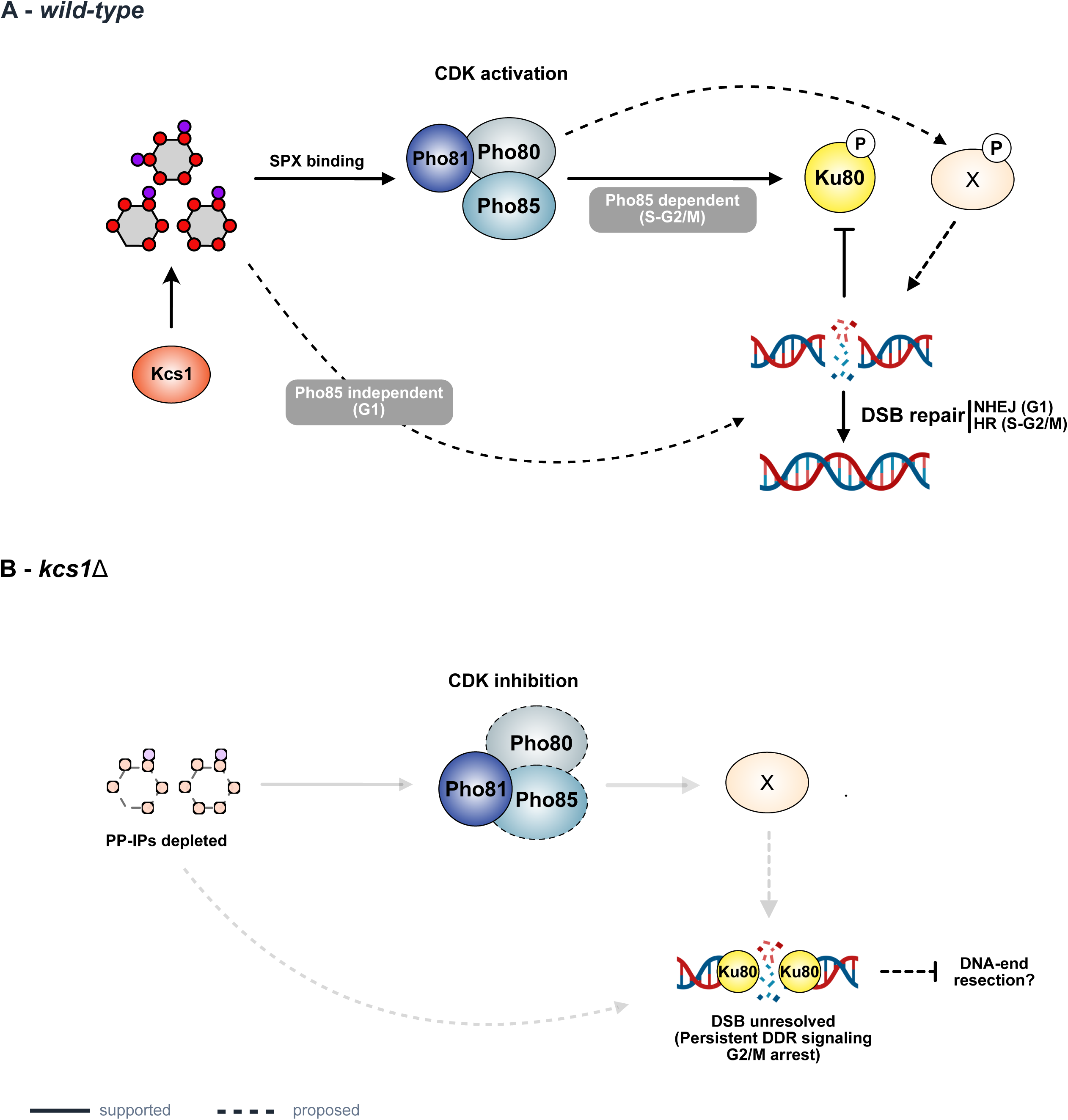
Working model for the Kcs1-Pho80-Pho85-Ku80 axis in DSB repair. (A) In wild-type cells, Kcs1-dependent inositol pyrophosphates (5-IP7 and/or 5-PP-IP4) accumulate upon genotoxic stress and bind the SPX domain of Pho81, relieving its inhibition of the Pho80-Pho85 cyclin-dependent kinase. Active Pho80-Pho85 regulates Ku80 and other proteins required for efficient double-strand break (DSB) repair. A parallel, Pho85-independent contribution of Kcs1-dependent PP-IPs to Ku80 regulation is also proposed. This axis operates independently of the core HR machinery and of phosphate homeostasis, and is active in G1, S, and G2/M. (B) Loss of Kcs1 depletes PP-IPs; Pho81, via its SPX domain, binds and inhibits the Pho80-Pho85 complex, leading to misregulation of Ku80 and other targets, failure to repair DSBs, persistent Rad53/γH2A signaling, G2/M arrest, and zeocin hypersensitivity. Dashed arrows and question marks denote steps that were inferred from genetic data and were not measured directly in this study, including the change in PP-IP levels upon genotoxic stress, the activity state of Pho80-Pho85, and the phosphorylation of Ku80.

Because both PP-IP metabolism and CDK control of DNA repair are conserved from yeast to humans, we expect analogous mechanisms to operate in other eukaryotes. In mammalian cells, IP6K inhibitors are being developed as metabolic and anticancer agents. Therefore, our results raise the possibility that part of their cellular activity reflects a compromised DDR rather than altered energy or phosphate metabolism alone. Also, in pathogenic fungi, if this pathway supports genome maintenance under host-imposed genotoxic stress, interfering with PP-IP synthesis could sensitize fungal pathogens to DNA-damaging insults.

## Experimental procedures

### Strains and plasmids

Unless otherwise stated, yeast strains used in this study were derived from S288C. A complete list of strains and plasmids is provided in Tables S1 and S2, respectively. Gene-deletion and epitope-tagged strains were constructed using a one-step gene replacement strategy (76, 77). Gene deletions were generated by replacing the endogenous allele with drug-resistance cassettes or auxotrophic markers amplified by PCR (78). Yeast transformations were performed using the lithium acetate method (79, 80). Correct integration of gene deletions was verified by PCR using locus-specific primers, and expression of epitope-tagged proteins was confirmed by western blotting. A detailed list of all the primers used in this study can be found in Table S3. To construct the Kcs1-3xFLAG strain, the *KCS1-3xFLAG* open reading frame, including a 50-bp upstream region, was cloned into the pFA6a-*NATMX6* vector using the SalI and AscI restriction sites. A 400-bp downstream region of *KCS1* was subsequently cloned into the plasmid using the PmeI and EcoRI restriction sites. The resulting *KCS1-3xFLAG* construct was excised with SalI and EcoRI and integrated at the *KCS1* locus of a strain in which *KCS1* had been replaced with the *HPMX6* cassette (*kcs1Δ::HPMX6*), which confers hygromycin resistance. Transformants able to grow on YPD containing clonNAT (nourseothricin), but not on YPD containing hygromycin, were selected. *KCS1-3xFLAG* expression was confirmed by western blotting. The K778A mutation was introduced by site-directed mutagenesis with mutagenic oligonucleotide primers following a protocol modified from (81). All plasmid constructs were sequence-verified by Plasmidsaurus Whole Plasmid Sequencing (Oxford Nanopore, R10.4.1).

### Cell growth, media, and synchronization

All strains were grown in YPD (1% yeast extract, 2% peptone, and 2% dextrose) medium at 30°C on a shaker at 250 rpm to logarithmic phase unless stated otherwise. For stable isotope labeling by amino acids in cell culture (SILAC)-based proteomic analyses, yeast cells were cultured in synthetic medium lacking arginine and lysine (CSM-Arg-Lys) and supplemented with either normal L-arginine and L-lysine (light culture) or ^13^C6,^15^N4 (heavy culture). Proline (80 mg/L) was added to prevent the metabolic conversion of arginine to proline. For synchronization, cells were treated either with α-factor (1 μg/mL) or nocodazole (10 μg/mL) for 2 hours followed by zeocin treatment (100 μg/mL) to induce DNA damage in synchronized yeast cultures. For Latrunculin A experiments, exponentially growing cells were treated with zeocin (50 µg/mL), Latrunculin A (20 µM in DMSO), or both, for 80 min at 30°C. Control cultures received an equivalent volume of DMSO (1%). Drug concentrations, incubation time and the γ-H2A readout were chosen to match those used by Hurst et al. (55), allowing direct comparison with the conditions under which nuclear actin was shown to drive INO80C-dependent conversion of zeocin-induced lesions into DSBs.

### Serial dilution and growth curve assays

Yeast strains of interest were cultured to the exponential phase and normalized to an optical density (OD600) of 1.0. Serial dilutions were then prepared in sterile 96-well plates, and the diluted cultures were transferred onto Petri dishes containing YPD agar supplemented with increasing concentrations of the indicated genotoxins using a metal replicator (Sigma-Aldrich). Cell growth was visually assessed after 48–72 h of incubation at 30°C, and images were acquired using a photodocumentation system (UVITEC 4.7, Cambridge).

For growth curve assays, yeast strains were cultured overnight in YPD medium until saturation. Cultures were subsequently diluted in YPD medium, with or without zeocin (10 μg/mL), to an initial OD600 of 0.1. Cell growth was monitored for 48 h at 30°C using a BioTek® Synergy H1 microplate reader. OD600 measurements were recorded at 10-minute intervals throughout the experiment, and growth curves were generated from the resulting measurements using GraphPad Prism version 11.0.1.

### Phosphate depletion from medium

Phosphate-depleted YPD medium was prepared by removing phosphate from yeast extract and peptone. Briefly, 20 g/L peptone and 10 g/L yeast extract were dissolved in distilled water in the presence of 2.46 g/L of MgSO4. With vigorous stirring, 4 mL of 25% ammonium hydroxide (NH₄OH) was added dropwise and the mixture was allowed to form a precipitate for 30 minutes at room temperature. The precipitated phosphate was initially removed by gravity filtration with Whatman #1 paper followed by vacuum filtration using a sterile Corning® bottle-top 0.22 μm vacuum filter system. The medium was adjusted to pH 6.5, autoclaved, and supplemented with 2% glucose after sterilization. Yeast cells were cultured in phosphate-depleted medium under the same conditions used for control cultures.

### Flow cytometry

Cell cycle analysis was performed as described before (46). Briefly, yeast cultures grown under specific conditions were collected, centrifuged, and the pellets were resuspended in 1 mL of 70% ethanol and incubated for 15 minutes at room temperature (or overnight at -20°C). After incubation, the fixed cells were centrifuged, and the pellet was dried using a speed vacuum. Then, sodium citrate buffer (50 mM, pH 7.2) was added, and the contents were sonicated (3x 3 s cycles, 35% amplitude) to break up the cell clusters. Samples were incubated with 100 µg of RNase A (QIAGEN^®^) for 1 hour at 37°C followed by incubation with 500 µg of Proteinase K (Invitrogen^®^) for 1 hour at 42°C. Finally, 1 µL of SYTOX Green (Life Technologies^®^) was added to the samples and analyzed using a flow cytometer (C6 Flow Cytometer BD^®^). Flow cytometry data were analyzed using FlowJo™ v10.8 Software (BD Life Sciences).

### Western blotting

Protein extracts were prepared by mechanical disruption with glass beads in lysis buffer containing 50 mM Tris-HCl (pH 7.5), 0.2% Tergitol, 150 mM NaCl, 5 mM EDTA, and protease and phosphatase inhibitor cocktails. Total protein extracts were denatured in SDS sample buffer supplemented with DTT. Proteins were separated by SDS-PAGE using 10% polyacrylamide gels for Rad53 analysis and 14% gels for γ-H2A detection. Enhanced resolution of Rad53 phospho-isoforms was performed as previously described (50). Proteins were transferred to polyvinylidene fluoride (PVDF) membranes (Amersham Hybond-P, GE Healthcare) at 100 V for 2 hours at 4°C. Membranes were probed with anti-γ-H2A (ab181447, Abcam; 1:4,000 dilution in Tris Buffered Saline with Tween (TBST) containing 5% nonfat dry milk), anti-Rad53 (ab104232, Abcam; 1:8,000 dilution in TBST containing 4% nonfat dry milk), and anti-Pgk1 (ab113687, Abcam; 1:15,000 dilution in TBST containing 2% nonfat dry milk), which was used as a loading control. Expression of epitope-tagged proteins was detected using the appropriate anti-tag antibodies. After primary antibody incubation, membranes were incubated with HRP-conjugated secondary antibodies (anti-mouse NA931 or anti-rabbit NA934, Cytiva) diluted 1:10,000 in TBST containing 2% nonfat dry milk. Signals were detected using either Clarity Max Western ECL Substrate (Bio-Rad) or Amersham ECL Prime (Cytiva), according to the manufacturers’ instructions. Chemiluminescent signals were acquired using either an Alliance 4.7 imaging system (UVITEC Cambridge, UK) or a ChemiDoc Imaging System (Bio-Rad Laboratories, USA).

### Inducible DSB reporter assay

The YRA213 reporter strain was used to assess cellular survival following induction of a site-specific DSB, as described in (82, 83). Briefly, a DSB is induced by the galactose-inducible HO endonuclease at a genomic *URA3* locus. For western blotting, the strains were inoculated in YPD medium overnight and diluted to an OD600 of 0.1 into YP medium containing 2% raffinose. Upon reaching the logarithmic phase, 2% galactose was added to induce HO endonuclease expression, and samples were subsequently collected at the indicated time points. Protein extraction and western blotting were performed as described above.

To measure cell survival following DSB induction, 12 independent colonies of each strain (wild-type and *kcs1Δ*) were inoculated into YPD medium and grown overnight. Cultures were then normalized to an OD600 of 1.0 and serially diluted. Aliquots of the diluted cultures were plated onto YPD agar plates as a non-inducing control and onto YPGal agar plates to induce HO expression. Colony formation was assessed after 72 h of incubation at 30°C.

### Fluorescence microscopy analysis

For the analysis of Rad52-mRuby2 foci, yeast cells were grown in YPD medium at 30°C until reaching mid-log phase. Zeocin was then added to a final concentration of 100 µg/mL, and cells were collected 2 hours after treatment. After centrifugation, cell pellets were resuspended in synthetic complete (SC) medium, and subsequent experimental procedures were performed as previously described (84). Live-cell imaging was performed using a Leica TCS SP8 confocal fluorescence microscope. For each biological replicate, at least 150 cells were scored and Rad52-mRuby2 foci were quantified from maximum-intensity projection images using LAS X Office software. Statistical analyses were performed as described in the Statistical analysis section.

### Mass spectrometry analysis

Yeast cells were cultured in 20 mL of SILAC light or heavy media until mid-logarithmic phase and treated with 100 μg/mL zeocin for 1 hour. Cell pellets were harvested, resuspended in TE buffer containing 1 mM Phenylmethylsulfonyl fluoride (PMSF) and phosphatase inhibitors, and stored at -80°C until processing. Proteins were extracted by bead beating in lysis buffer supplemented with protease and phosphatase inhibitors (50 mM Tris-HCl pH 8, 0.2% Tergitol; 150 mM NaCl; 5 mM EDTA; 1 mM PMSF; 5 mM sodium fluoride (NaF); Pierce Protease Inhibitor (Thermo Fisher)), and protein concentration was determined using the Bradford assay. Equal amounts of light and heavy-labeled proteins were mixed, denatured in 1% SDS, reduced with 5 mM DTT, alkylated with 15 mM iodoacetamide, precipitated with 50% acetone, 49.9% ethanol, and 0.1% acetic acid, resuspended in 8 M urea in 100 mM Tris-HCl (pH 8.0), and diluted with 50 mM Tris and 150 mM NaCl to a final urea concentration of 2 M. Samples were digested with mass spectrometry-grade modified trypsin (Trypsin Gold, Promega), desalted using Sep-Pak C18 cartridges (Waters), prefractionated by hydrophilic interaction liquid chromatography (HILIC) using an HPLC system (Dionex/Thermo Fisher Ultimate 3000), and analyzed by LC–MS/MS using a ThermoFisher Q Exactive HF mass spectrometer coupled to an Ultimate 3000 HPLC system. The resulting datasets were processed using Trans-Proteomics Pipeline with the Comet search engine (85) against the *Saccharomyces cerevisiae* protein database. The output was subsequently processed, statistically analyzed, and visualized using R (R Foundation for Statistical Computing).

### Functional enrichment analysis

Enrichment was performed in STRING v12.5 using the Proteins with Values/Ranks mode against local network clusters, with all 2,674 quantified proteins ranked by the mean log2 ratio of the two reciprocal SILAC labeling experiments. Significance was determined by the Kolmogorov– Smirnov test implemented in STRING with Benjamini–Hochberg correction, and clusters with FDR ≤ 0.05 were retained. Because STRING local network clusters are organized hierarchically, clusters sharing ≥ 70% of their member proteins with a more significant cluster were merged and are represented by the most significant member. The full list of clusters is given in Table S4. For the network in Fig. 5*C*, STRING associations were performed at medium confidence (combined score ≥ 0.400).

### Statistical analysis

Statistical analyses were performed using GraphPad Prism version 11.0.1 (GraphPad Software, USA). Unless otherwise stated, data are presented as mean ± SEM from independent biological replicates. The exact n for each experiment is given in the corresponding figure legend. Rad52 focus data (Fig. 4*D* and Fig. S4) were analyzed by two-way repeated-measures ANOVA matched on both factors (genotype and treatment) with Sidak’s correction for multiple comparisons. Survival after HO induction (Fig. 3*G*) was compared by an unpaired two-tailed t test. Growth curves are presented as the median of two biological replicates and were not tested statistically. P values below 0.05 were considered significant.

## Data Availability

The mass spectrometry proteomics data have been deposited to the ProteomeXchange Consortium via the PRIDE partner repository with the dataset identifier PXD084543. All other data are contained within the manuscript and its supporting information. Yeast strains and plasmids generated in this study (Tables S1 and S2) are available upon request.

## Supporting information

This article contains supporting information.

## Acknowledgments

We would like to thank Dr. Luis Eduardo Soares Netto for donating yeast strains and for allowing access to their laboratory and equipment to perform growth curve experiments. We also thank Dr. Francisco Meirelles Bastos de Oliveira (UFRJ) and Dr. Claudio Akio Masuda (UFRJ) for comments and suggestions, and Jacilene Barbosa and Dr. Gleice da Graça Rocha for technical support.

## Author contributions

J. R. R. C. conceptualization; M. M. W., M. B. S., and J. R. R. C. data curation; J. R. R. C., G. M. P., and M. M. W. formal analysis; J. R. R. C. and G. M. P. funding acquisition; J. R. R. C., G. M. P., P. P. S., G. M., and M. M. W. investigation; J. R. R. C., G. M. P., and M. B. S. methodology; J. R. R. C. project administration; J. R. R. C. and M. B. S. resources; J. R. R. C. supervision; G. M. P. and J. R. R. C. visualization; J. R. R. C. and G. M. P. writing– original draft; J. R. R. C., G. M. P., and M. B. S. writing–review & editing.

## Funding and additional information

This work was supported by the São Paulo Research Foundation (FAPESP) grants 2018/05417-0 and 2024/10865-3 (to J. R. R. C.). G. M. P. was supported by FAPESP doctoral (2021/04887-6) and BEPE research internship abroad (2023/09910-1) fellowships.

## Conflict of interest

The authors declare that they have no conflicts of interest with the contents of this article

## Abbreviations

1-IP7: 1-diphosphoinositol pentakisphosphate
5-IP7: 5-diphosphoinositol pentakisphosphate
1,5-IP8: 1,5-bis-diphosphoinositol tetrakisphosphate
CDK: cyclin-dependent kinase
CKI: CDK inhibitor
DDR: DNA damage response
DSB: double-strand break
FDR: false discovery rate
HO: homothallic switching endonuclease
HR: homologous recombination
HU: hydroxyurea
IP: inositol polyphosphate
IP3: inositol 1,4,5-trisphosphate
IP6: inositol hexakisphosphate
IPK: inositol polyphosphate kinase
MEF: mouse embryonic fibroblast
MMS: methyl methanesulfonate
NHEJ: non-homologous end joining
Pi: inorganic phosphate
PIP2: phosphatidylinositol 4,5-bisphosphate
polyP: inorganic polyphosphate
PP-IP: inositol pyrophosphate
PP-IPk: inositol pyrophosphate kinase
SILAC: stable isotope labeling by amino acids in cell culture
SPX: SYG1/Pho81/XPR1 domain
YPD: yeast extract–peptone–dextrose medium
SC: synthetic complete medium;
CSM: complete supplement mixture medium.

## Notes

### Competing Interest Statement

The authors have declared no competing interest.

### Summary of Updates

This version was reformatted according to journal style. The Author Summary was removed. Sections were reordered and renamed, and the citation style for references, figures and supplementary items was updated. A list of abbreviations and a statement on reagent availability were added. The Author contributions section was revised, and minor grammatical and editorial corrections were made throughout. The data, analyses and conclusions are unchanged.

