## Supporting Information for "Kcs1-dependent inositol pyrophosphates promote DNA double-strand break repair via Pho80-Pho85 in budding yeast"

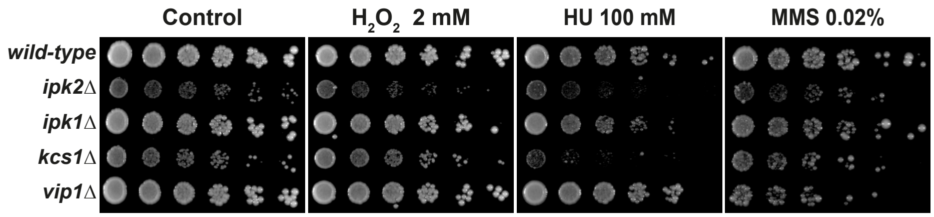


**Figure S1. Genetic interactions between *KCS1* and other inositol polyphosphate kinases across genotoxic agents.** Five-fold serial dilutions of wild-type, *ipk2Δ*, *ipk1Δ*, *kcs1Δ* and *vip1Δ* cells were spotted onto YPD plates containing $H_2O_2$ (2 mM), hydroxyurea (100 mM), MMS (0.02%), or onto drug-free YPD (Control).

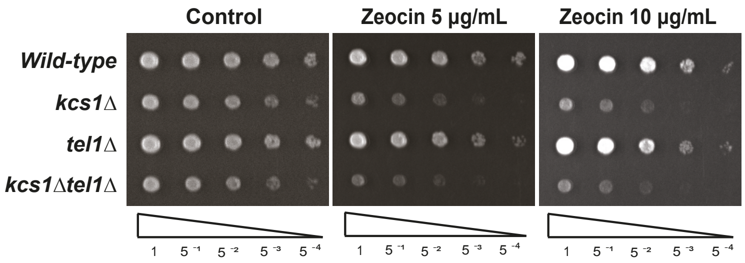


**Figure S2. Deletion of *TEL1* does not suppress the zeocin sensitivity of *kcs1Δ* cells.** Five-fold serial dilutions of wild-type, *kcs1Δ*, *tel1Δ* and *kcs1Δtel1Δ* cells were spotted onto YPD plates containing the indicated concentrations of zeocin.

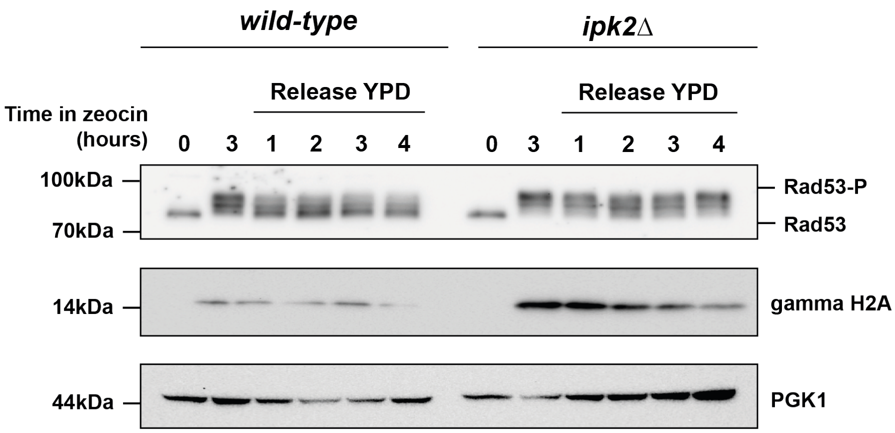


**Figure S3.** ***ipk2Δ*** **cells sustain DNA damage signaling during recovery from zeocin treatment.** Exponentially growing wild-type and *ipk2Δ* cells were treated with zeocin (100 µg/mL) for 3 h, washed, and released into drug-free YPD (Release YPD). Samples were collected before treatment (0), after 3 h in zeocin, and 1, 2, 3 and 4 h after release. Rad53 phosphorylation (Rad53-P), γ-H2A and Pgk1 (loading control) were analyzed by western blotting; molecular mass markers are indicated on the left. Blots are representative of two independent experiments.

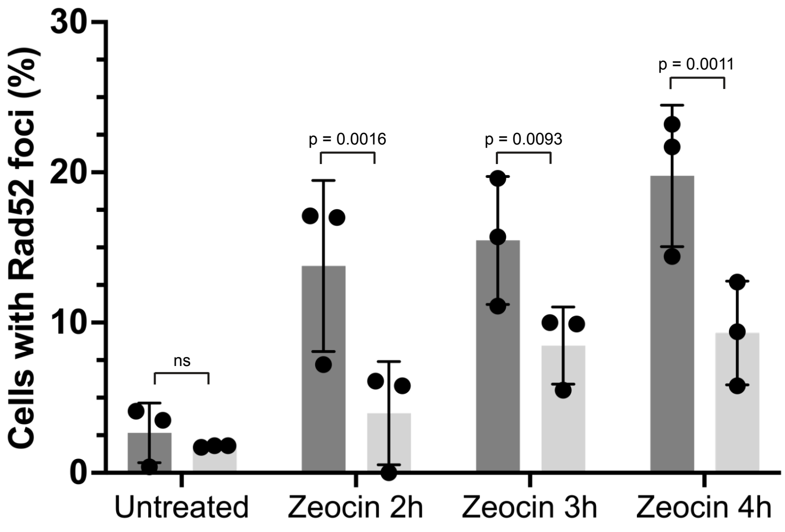


**Figure S4. Rad52 focus formation is reduced in asynchronous *kcs1Δ* cultures.** Quantification of Rad52-mRuby2 nuclear foci in asynchronously growing wild-type (dark bars) and *kcs1Δ* (light bars) cells, left untreated or treated with zeocin (100 µg/mL) for 2, 3 or 4 h. Bars represent the mean percentage of cells containing at least one Rad52-mRuby2 focus from three independent biological replicates; individual replicate values are shown as symbols and error bars represent SEM. At least 150 cells were scored per strain and condition in each replicate. Statistical analysis was performed by two-way ANOVA matched on both factors (genotype and treatment), with Sidak's correction for multiple comparisons applied to genotype comparisons within each treatment and using separate variances.

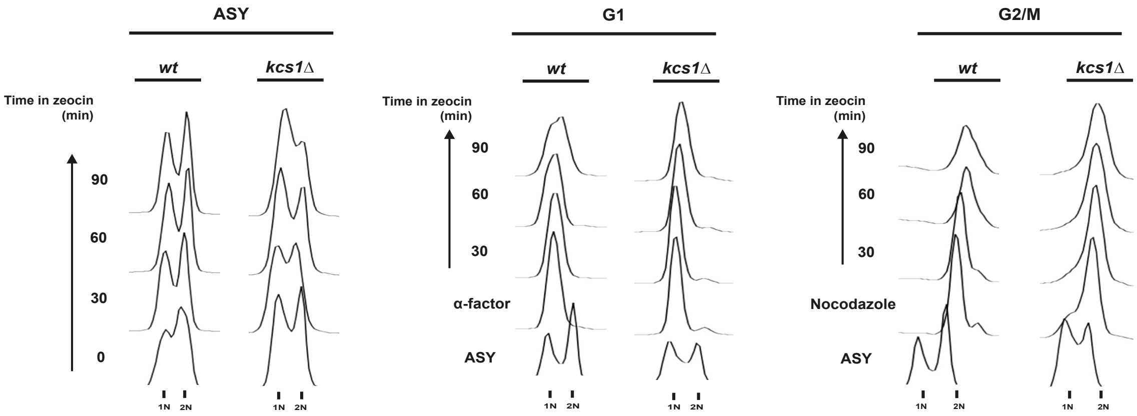


**Figure S5. Cell-cycle synchronization of wild-type and *kcs1Δ* cells during zeocin treatment.** Flow cytometry profiles of DNA content (1N and 2N positions indicated) for wild-type (wt) and *kcs1Δ* cells. Left, asynchronous (ASY) cultures treated with zeocin (100 µg/mL) for 0, 30, 60 and 90 min. Middle, cells arrested in G1 with α-factor (1 µg/mL) before zeocin addition. Right, cells arrested in G2/M with nocodazole (10 µg/mL) before zeocin addition. Both strains were uniformly arrested at the intended stage and remained arrested throughout treatment. Profiles are representative of three independent experiments.

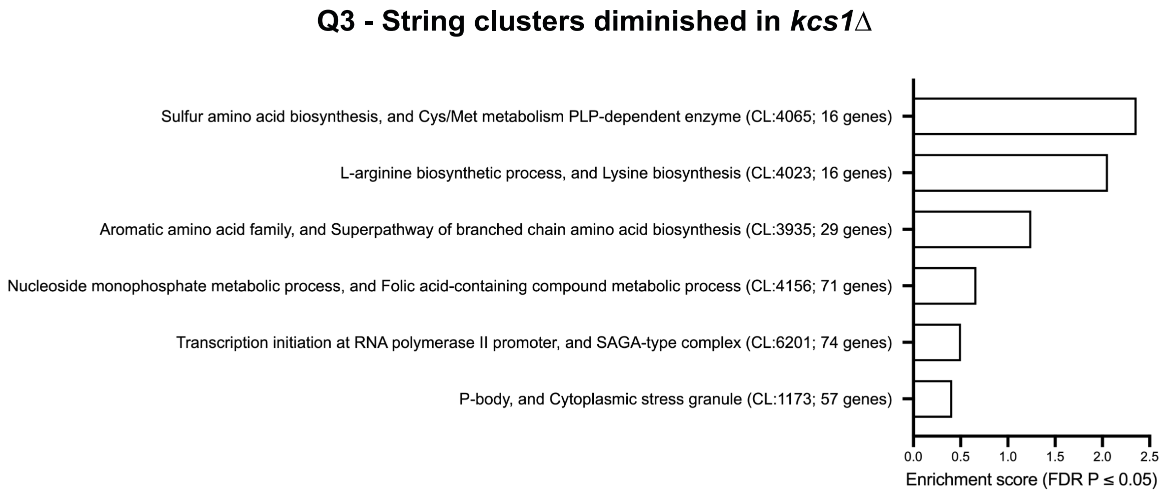


**Figure S6. Functional modules enriched among proteins decreased in *kcs1Δ* cells.** Rank-based enrichment of all 2,674 quantified proteins against STRING local network clusters. STRING cluster identifiers and the number of member proteins quantified here are given in parentheses.

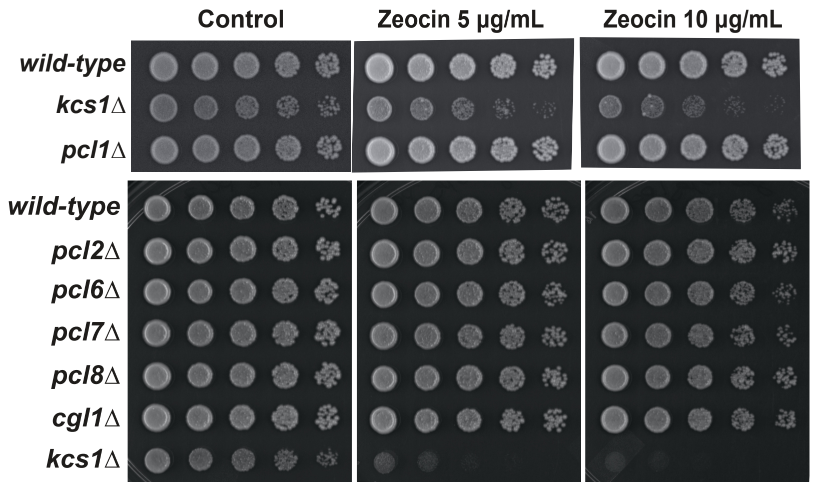


**Figure S7. Deletion of Pho85 cyclins other than Pho80 does not confer zeocin sensitivity.** (A) Five-fold serial dilutions of wild-type, *kcs1Δ* and *pcl1Δ* cells and (B) of wild-type, *pcl2Δ*, *pcl6Δ*, *pcl7Δ*, *pcl8Δ*, *cgl1Δ* and *kcs1Δ* cells were spotted onto YPD plates containing zeocin or onto drug-free YPD (control) and incubated at 30°C for 2–3 days. *kcs1Δ* cells are included in both panels as a sensitivity control.

**Table S1.** Yeast strains used in this study. This table is provided as a separate excel file titled “Table S1 strains”

**Table S2.** Plasmids used in this study. This table is provided as a separate excel file titled “Table S2 plasmids”

**Table S3.** Primers used in this study. This table is provided as a separate excel file titled “Table S3 primers”

**Table S4.** Functional enrichment with STRING local network clusters. This table is provided as a separate excel file titled “Table S4 STRING clusters”

**File S1.** SILAC proteomics dataset (wild-type vs *kcs1Δ*, zeocin-treated). This table is provided as a separate excel file titled “S1 File”
